# Mechanochemical control of epidermal stem cell divisions by B-plexins

**DOI:** 10.1101/2020.04.30.070359

**Authors:** Chen Jiang, Ahsan Javed, Laura Kaiser, Michele M. Nava, Dandan Zhao, Dominique T. Brandt, Javier Fernández-Baldovinos, Luping Zhou, Carsten Höß, Kovilen Sawmynaden, Arkadiusz Oleksy, David Matthews, Lee S. Weinstein, Hermann-Josef Gröne, Carien M. Niessen, Stefan Offermanns, Sara A. Wickström, Thomas Worzfeld

## Abstract

The precise spatiotemporal control of cell proliferation is key to the morphogenesis of epithelial tissues. Epithelial cell divisions lead to tissue crowding and local changes in force distribution, which in turn suppress the rate of cell divisions. However, the molecular mechanisms underlying this mechanical feedback are largely unclear. Here, we identify a critical requirement of B-plexin transmembrane receptors in the response to crowding-induced mechanical forces during embryonic skin development. Epidermal stem cells lacking B-plexins fail to sense mechanical compression, resulting in disinhibition of the transcriptional coactivator YAP, hyperproliferation, and tissue overgrowth. Mechanistically, we show that B-plexins mediate mechanoresponses to crowding through stabilization of adhesive cell junctions and lowering of cortical stiffness. Finally, we provide evidence that the B-plexin-dependent mechanochemical feedback is also pathophysiologically relevant to limit tumor growth in basal cell carcinoma, the most common type of skin cancer. Our data uncover a central role of B-plexins in mechanosensation to couple cell density and cell division in development and disease.

## INTRODUCTION

During embryogenesis, tissue growth and shape are intimately connected^1^. Tissue growth intrinsically generates mechanical strains and compressions, and, conversely, mechanical deformations provide a fundamental regulatory feedback for growth control^1–3^. This feedback requires cells to sense mechanical forces, which are then transformed into biochemical signals that in turn control cellular functions including the rate of cell divisions^3–6^. Compression of cells within a tissue results in suppression of proliferation, a response termed contact inhibition of proliferation^2, 7^. In the mammalian embryonic epidermis, proliferation of stem cells within the mechanically jammed basal layer causes crowding and local cell stress anisotropy^8–10^. How epidermal stem cells sense these mechanical forces and couple them with division is largely unknown.

Plexins constitute a family of single-pass transmembrane proteins that have initially been described for their role as axon guidance receptors for semaphorins in the developing nervous system^11, 12^. In vertebrates, nine plexins have been identified, which – based on homology – are grouped into four subfamilies, Plexin-A1-4, Plexin-B1-3, Plexin-C1, and Plexin-D1^13^ They are now recognized to be of central importance for cell-cell communication in multiple tissues and biological systems, such as the immune and bone system, as well as in cancer^14–16^. Very recently, Plexin-D1 has been shown to be activated by fluid shear stress-induced mechanical forces in vascular endothelial cells^17^. Whether plexins play a role in mechanosensation in epithelial cells is unknown.

Here, we report that epidermal stem cells require Plexin-B1 and Plexin-B2 to sense mechanical compression, and uncover Plexin-B1 and Plexin-B2 as negative upstream regulators of YAP that suppress stem cell divisions in response to crowding. This biomechanical feedback mechanism ensures a robust control of proliferation rates by cell density, and is functionally relevant for both tissue morphogenesis as well as cancer.

## RESULTS

### Plexin-B1/Plexin-B2 suppress epidermal stem cell proliferation during embryonic development

Divisions of stem cells within an epithelial layer induce crowding and cell shape anisotropy^18^. We hypothesized that a mechanical feedback to control stem cell divisions would therefore be particularly relevant during developmental stages with high stem cell proliferation rates. We observed that in the developing mouse epidermis, stem cell proliferation rates were highest at embryonic day 15.5 (E15.5) and gradually declined to very low rates in the adult (Fig. 1a,b), consistent with previous reports^19, 20^. The mRNA expression levels of plexins showed a striking positive correlation with proliferation rates, with Plexin-B2 being the most highly expressed plexin family member during embryonic development (Fig. 1c and Supplementary Fig. 1a,b). Immunostainings revealed that Plexin-B2 was expressed in all layers of the developing epidermis including the K14-positive stem cells (Fig. 1d,e). Plexin-B1, a receptor highly homologous to Plexin-B2^11, 13^, localized exclusively to the stem cell layer (Fig. 1d). To identify the functional role of Plexin-B1 and Plexin-B2 in epidermal stem cell proliferation, we inactivated the respective genes. Given the overlapping expression pattern of Plexin-B1 and Plexin-B2 in K14-positive stem cells (Fig. 1d) and the functional redundancy of these two receptors during development^21–23^, we generated mice lacking both Plexin-B1 and Plexin-B2 in epidermal stem cells by crossing conditional alleles for Plexin-B1 and Plexin-B2 with a constitutively expressed keratin 14-Cre line (K14-Cre;*plxnb1*^flox/flox^;*plxnb2*^flox/flox^). Consistent with the onset of Cre expression^24^, Plexin-B1 and Plexin-B2 expression was lost as early as E15.5 (Supplementary Fig. 1c). Interestingly, Plexin-B1/Plexin-B2 double-knockout mice (“PlexDKO”) showed a marked epidermal hyperproliferation (Fig. 1f), resulting in epidermal thickening (Fig. 1g,h). This epidermal thickening was due to an expansion of K14-positive stem cells, while the number of K10-positive differentiated cells was not significantly increased (Fig. 1i,j and Supplementary 1d). Apoptosis rates remained unchanged (Supplementary Fig. 1e). Taken together, these results indicate that Plexin-B1 and Plexin-B2 suppress epidermal stem cell proliferation during embryonic development.

**Fig. 1:**
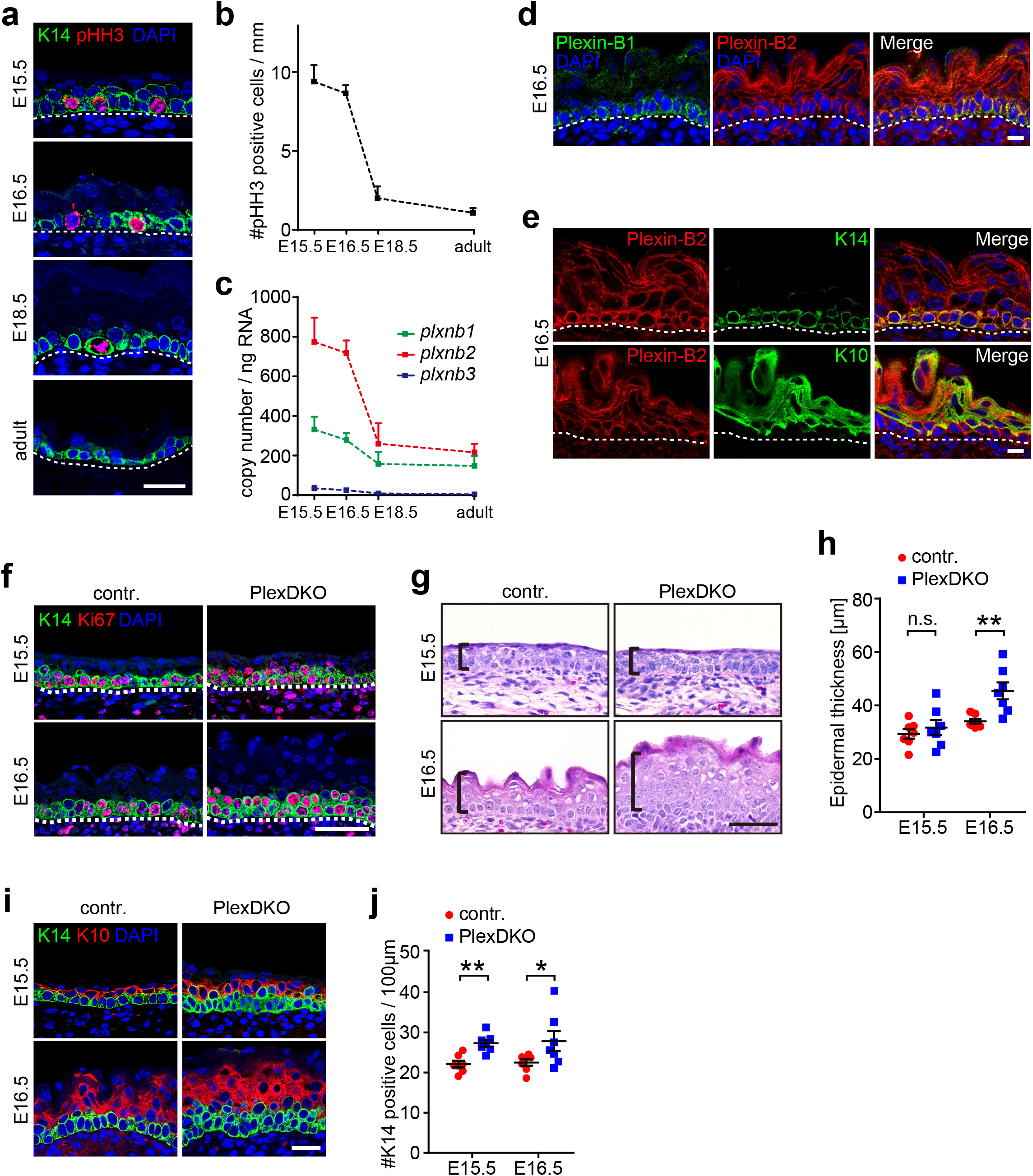
Plexin-B1/Plexin-B2 suppress epidermal stem cell proliferation during embryonic development. **a**, Confocal images of immunostainings of murine skin at embryonic day 15.5 (E15.5), 16.5 (E16.5), 18.5 (E18.5) and at an adult stage using anti-phospho-histone H3 (pHH3; red) and anti-keratin 14 (K14; green) antibodies. Dashed lines indicate the basement membrane. Scale bar, 25 μm. **b**, Quantification of pHH3-positive cells at different time points of embryonic (E) development and in the adult (mean ± s.e.m.; *n*=3 mice per time point). **c**, mRNA expression levels of genes encoding B-plexins (*plxnb*) in the murine skin at the indicated time points as determined by quantitative RT-PCR (mean ± s.e.m.; *n*=3 mice per time point). **d**, Confocal images of immunostainings of murine epidermis at embryonic day 16.5 (E16.5) using anti-Plexin-B1 (green) and anti-Plexin-B2 antibodies (red). Dashed lines indicate the basement membrane. Scale bar, 10 μm. **e**, Confocal images of immunostainings of murine epidermis at E16.5 using anti-Plexin-B2 (red), and anti-keratin 14 (K14) or anti-keratin 10 (K10) antibodies (green). Scale bar, 10 μm. **f**, Confocal images of immunostainings of murine epidermis at embryonic day 15.5 and 16.5 using an anti-Ki67 (red) and anti-keratin 14 (green) antibodies. “contr.”: control mice (genotype *plxnb1*^flox/flox^;*plxnb2*^flox/flox^), “PlexDKO”: epidermisspecific Plexin-B1/Plexin-B2 double-knockout mice (genotype K14-Cre;*plxnb1*^flox/flox^;*plxnb2*^flox/flox^). Blue: DAPI. Scale bar, 50 μm. **g**, H&E stained histological sections of murine epidermis at the indicated time points of embryonic development. Brackets indicate epidermal thickness. Scale bar, 50 μm. **h**, Quantification of the data in (g) (mean ± s.e.m.; E15.5 and E16.5: *n*=7 mice per genotype and time point; unpaired t-test). **i**, Confocal images of immunostainings of murine embryonic epidermis using anti-K14 (green) and anti-K10 (red) antibodies. Scale bar, 25 μm. **j**, Quantification of K14-positive cells (mean ± s.e.m.; *n*=7 mice per genotype and time point; unpaired t-test).

### Hyperproliferation of Plexin-B1/Plexin-B2 double-deficient epidermal stem cells induces overcrowding, cell shape anisotropy and differentiation

We next investigated how the increased number of epidermal stem cell divisions in Plexin-B1/Plexin-B2 double-knockout mice affected stem cell density and shape. To do so, we performed segmentation analyses of the epidermal stem cell layer at embryonic day 15.5 (Fig. 2a). In mice lacking Plexin-B1 and Plexin-B2, we observed both a higher cell density (Fig. 2b) as well as smaller and more variable cell sizes (Fig. 2c) than in control mice. This was accompanied by an increase in cell shape anisotropy, with cells being more elongated (Fig. 2d,e). Similar abnormalities were detected at embryonic day 16.5 (Supplementary Fig. 2a-e). These data demonstrate that Plexin-B1/Plexin-B2 double-deficiency results in overcrowding and shape deformation in the epidermal stem cell layer.

**Fig. 2:**
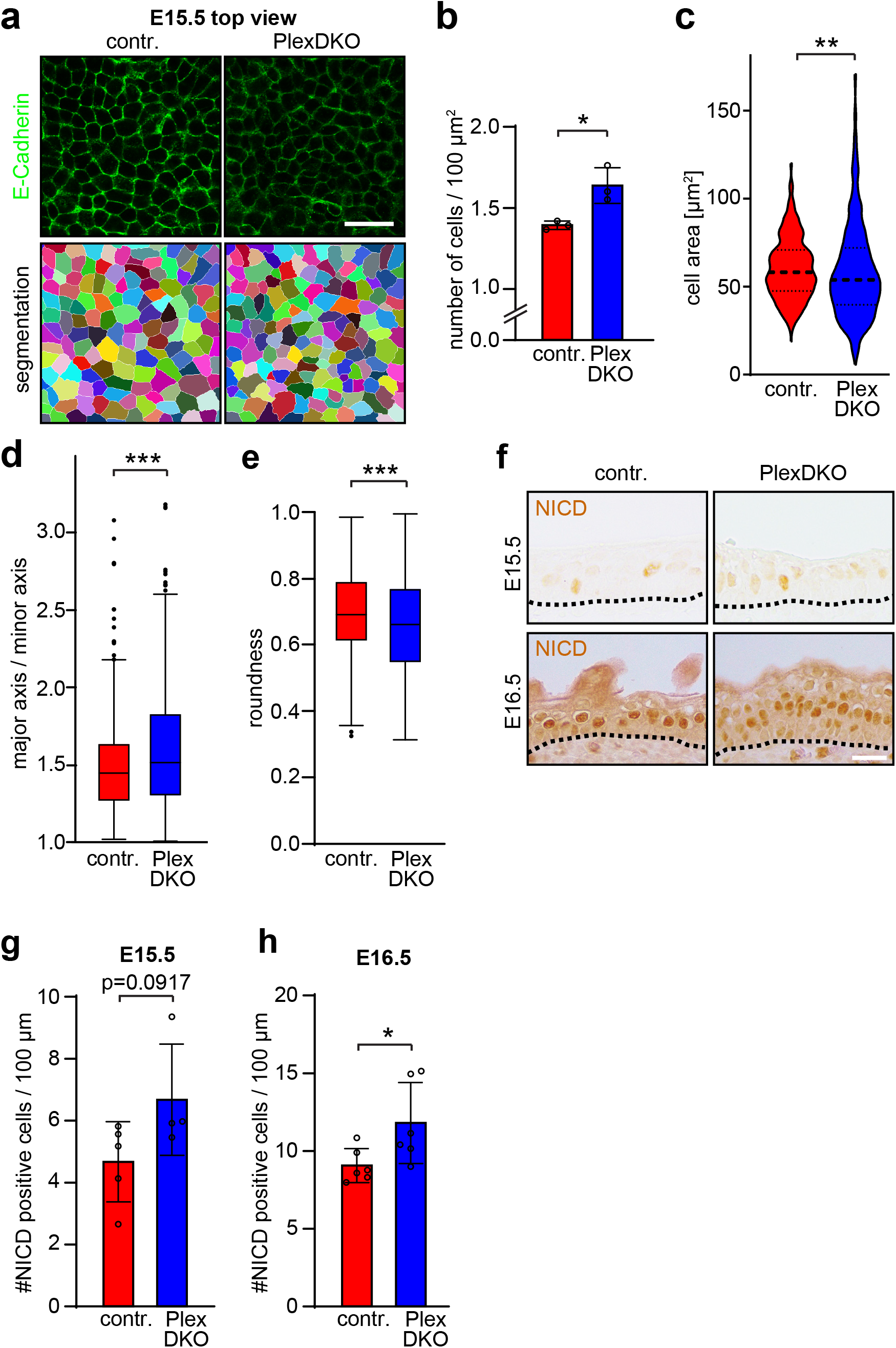
Hyperproliferation of Plexin-B1/Plexin-B2 double-deficient epidermal stem cells induces overcrowding, cell shape anisotropy and differentiation. **a**, Upper row: Confocal images of whole-mount immunostainings (top view) of murine epidermis at E15.5 using an anti-E-cadherin antibody (green). Shown are representative images of the basal cell layer. Lower row: Segmentation analyses of the images depicted in the upper row. Scale bar, 25 μm. **b**, Quantification of the number of epidermal stem cells (basal layer) per area (mean ± s.d.; *n*=3 mice per genotype). **c**, Analysis of epidermal stem cell areas (violin plot with first quartile, median, and third quartile; control: *n*=415 from 3 mice, PlexDKO: *n*=509 from 3 mice; F-test of equality of variances: p<0.0001, Mann-Whitney U test: p=0.0011). **d,e,** Quantification of cell shape anisotropy (box plot with minimum, first quartile, median, third quartile and maximum; control: *n*=415 from 3 mice, PlexDKO: *n*=509 from 3 mice; Mann-Whitney U test: p=0.0003). **f**, Immunohistochemistry on murine embryonic epidermis using an anti-NICD antibody (brown). Dashed lines indicate the basement membrane. Scale bar, 25 μm. **g,h**, Quantification of the data in (f) (mean ± s.d.; E15.5 control: *n*=5 mice, E15.5 PlexDKO: *n*=4 mice, E16.5: *n*=6 mice per genotype; unpaired t-test).

It is known that proliferation-induced crowding and cell shape distortion trigger differentiation and delamination of epidermal stem cells^18^. To assess whether loss of Plexin-B1/Plexin-B2 affects stem cell differentiation, we performed immunostainings for active Notch (Notch intracellular domain = NICD), which plays a decisive role in epidermal differentiation and specifically labels epidermal cells committed to differentiation^25^. While the number of NICD-positive cells in the Plexin-B1/Plexin-B2-deficient epidermis was not significantly changed at embryonic day 15.5 (Fig. 2f,g) – the developmental stage when stem cell hyperproliferation was most pronounced (Fig. 1f) –, Notch signaling became markedly increased at embryonic day 16.5 (Fig. 2f,h). We conclude that hyperproliferation of Plexin-B1/Plexin-B2-deficient epidermal stem cells coincides with overcrowding and cell shape anisotropy in the basal layer, which is followed – with a temporal delay – by increased differentiation to restore tissue size and architecture.

### Plexin-B1/Plexin-B2 inhibit YAP activity in response to mechanical forces

We next sought to identify the downstream signaling mechanism by which Plexin-B1/Plexin-B2 balance cell division rates to cell density. The transcriptional regulator YAP is known as a key factor in sensing cell density and shape to regulate cell proliferation^7, 26^. In the epidermis, high cell densities suppress YAP activity and stem cell divisions^27, 28^. We therefore examined how stem cell overcrowding and shape anisotropy in the Plexin-B1/Plexin-B2-deficient embryonic epidermis impact on YAP signaling. Surprisingly, we observed that both the levels of active, dephosphorylated YAP (Fig. 3a-c) as well as the mRNA expression level of the YAP target gene *ctgf* (Supplementary Fig. 3a-c) were elevated in embryonic epidermal stem cells lacking Plexin-B1 and Plexin-B2. This suggested that Plexin-B1/Plexin-B2-deficient epidermal stem cells fail to restrict YAP activity in response to crowding. To test this hypothesis, we cultured primary mouse keratinocytes at different densities and analyzed YAP localization as well as mRNA expression levels of YAP target genes. At low cell densities, when YAP is predominantly localized to the nucleus and active, these parameters were uninfluenced by the loss of Plexin-B1/Plexin-B2 (Supplementary Fig 3d,e). At high cell densities, however, Plexin-B1/Plexin-B2-deficiency impeded shuttling of YAP out of the nucleus (Supplementary Fig. 3h) and resulted in elevated mRNA expression levels of YAP target genes (Supplementary Fig. 3i). Interestingly, the suppressive effect of Plexin-B1/Plexin-B2 on YAP activity at high cell density was dependent on the presence of calcium in the culture medium (Supplementary Fig. 3f-i), which induces the formation of cadherin-based cell-cell adhesion complexes^29^. To further assess the role of Plexin-B1/Plexin-B2 in mechanosensation of tissue stress anisotropy caused by crowding, we seeded primary mouse keratinocytes on circular or square micropatterned surfaces with identical surface areas, on which cells experience isotropic (circles) and anisotropic (squares) traction forces^18^. Fully in line with a function of Plexin-B1/Plexin-B2 in mechanosensation of crowding, the dependence of YAP localization on Plexin-B1/Plexin-B2 was more pronounced under anisotropic than under isotropic traction stress conditions (Fig. 3d,e). Finally, to directly test for a requirement of Plexin-B1/Plexin-B2 in mechanosensation of crowding-induced mechanical forces, we subjected primary mouse keratinocyte monolayers to static stretch (for 12 hours) to expand the cell-substrate adhesive surface area, followed by release of tension resulting in a transient increase in monolayer compression^18^ (Fig. 3f). While in control cells, compression triggered exclusion of YAP from the nucleus, Plexin-B1/Plexin-B2-deficient cells failed to respond (Fig. 3g,h). Taken together, these data indicate that Plexin-B1/Plexin-B2 are required for mechanosensation in epidermal stem cells, and suppress YAP activity in response to mechanical forces generated by crowding-induced lateral compression.

**Fig. 3:**
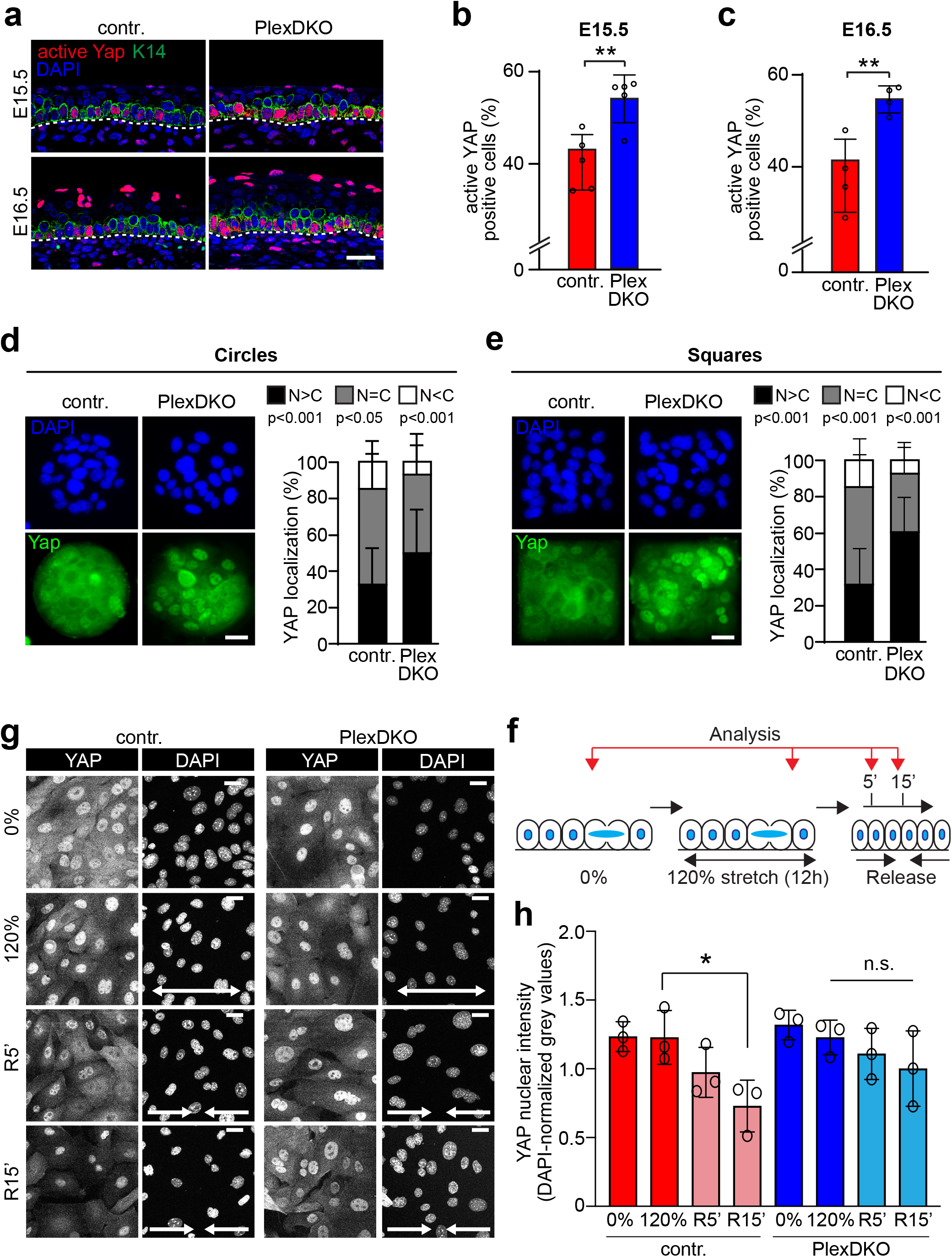
Plexin-B1/Plexin-B2 inhibit YAP activity in response to mechanical forces. **a**, Confocal images of immunostainings of murine epidermis at E15.5 (upper row) and E16.5 (lower row) using anti-active YAP (red) and anti-K14 antibodies (green). Scale bar, 25 μm. **b,c**, Quantification of the data in (a), i.e. of cells positive for active Yap (percentage of basal cells) (mean ± s.d.; E15.5: *n*=5 mice per genotype, E16.5: *n*=4 mice per genotype; unpaired t-test). **d,e**, Primary mouse keratinocytes on (d) circular or (e) square micropatterns immunostained for Yap (green) after 3 h of 1.8 mM Ca^2+^. Representative images are shown on the left, quantifications of Yap localization are shown on the right (mean ± s.d.; *circles*: control: *n*=73 micropatterns from 5 mice, PlexDKO: *n*=73 micropatterns from 4 mice; N>C: p=0.000008, N=C: p=0.012038, N<C: p=0.000008; *squares*: control: *n*=69 micropatterns from 5 mice, PlexDKO: *n*=69 micropatterns from 4 mice; N>C: p<0.000001, N=C: p<0.000001, N<C: p=0.000050; unpaired t-test). Blue: DAPI. Scale bars, 25 μm. **f**, Schematic illustration of abrupt crowding experiments. Confluent monolayers were exposed to 120% static uniaxial stretch, which was abruptly released. Cells were analyzed at the indicated time points (R5’: 5 minutes after release; R15’: 15 minutes after release). **g,h**, Representative YAP immunofluorescence images (g) and quantification of nuclear YAP (h) upon crowding (mean ± s.d.; *n*=3 independent experiments with >200 cells/condition/experiment; *p=0.0124, Friedman/Dunn’s; scale bars 20 μm).

### Plexin-B1/Plexin-B2 mediate mechanosensation through stabilization of adhesive cell-cell junctions

We next aimed at the elucidation of the precise molecular mechanism through which Plexin-B1/Plexin-B2 limit mechanosensitive YAP activity and epidermal stem cell proliferation. Plexin-B1 and Plexin-B2 are known to serve as receptors for class-4 transmembrane semaphorins, and to mediate forward signaling in response to ligand binding^13^. In addition, both Plexin-B1 as well as Plexin-B2 can act as ligands and induce reverse signaling via binding to class-4 transmembrane semaphorins^30, 31^. To investigate the effect of an activation of semaphorin-plexin signaling on YAP activity, we treated primary mouse keratinocytes with recombinant semaphorin 4C (Sema4C), a ligand for Plexin-B2, or with recombinant Plexin-B1 or Plexin-B2, and measured the mRNA expression levels of the YAP target genes *ctgf* and *cyr61*. However, neither activation of semaphorin-plexin forward nor of reverse signaling was sufficient to suppress YAP target gene expression (Supplementary Fig. 4a). Interestingly, in Xenopus, Plexin-A1 has been described to interact in *trans* with Plexin-A1 on other cells, and this homophilic binding has been shown to depend on the presence of calcium ions^32^. We found that the localization of Plexin-B1 and Plexin-B2 to cell-cell contacts of primary mouse keratinocytes was stabilized in the presence of calcium (Fig. 4a and Supplementary Fig. 4b), which was intriguing given the strong calcium-dependency of YAP activity control by Plexin-B1/Plexin-B2 (Supplementary Fig. 3d-i). Structure-function experiments in a renal tubular epithelial cell line showed that this localization of Plexin-B2 to cell-cell contacts relied on its extracellular domain and was independent of its intracellular domain (Supplementary Fig. 4c). To test whether Plexin-B1 and Plexin-B2 mediate homophilic binding in the presence of calcium, we measured the adhesion of primary mouse keratinocytes to recombinant Plexin-B1 and Plexin-B2. Indeed, while control cells strongly adhered to recombinant Plexin-B1 and Plexin-B2, Plexin-B1/Plexin-B2-deficient cells adhered less efficiently (Fig. 4b). In contrast, we did not detect any binding of primary mouse keratinocytes to recombinant Sema4C (Supplementary Fig. 4d). These data show that Plexin-B1/Plexin-B2 localize to cell junctions and promote cell-cell adhesion via a homophilic binding mechanism.

**Fig. 4:**
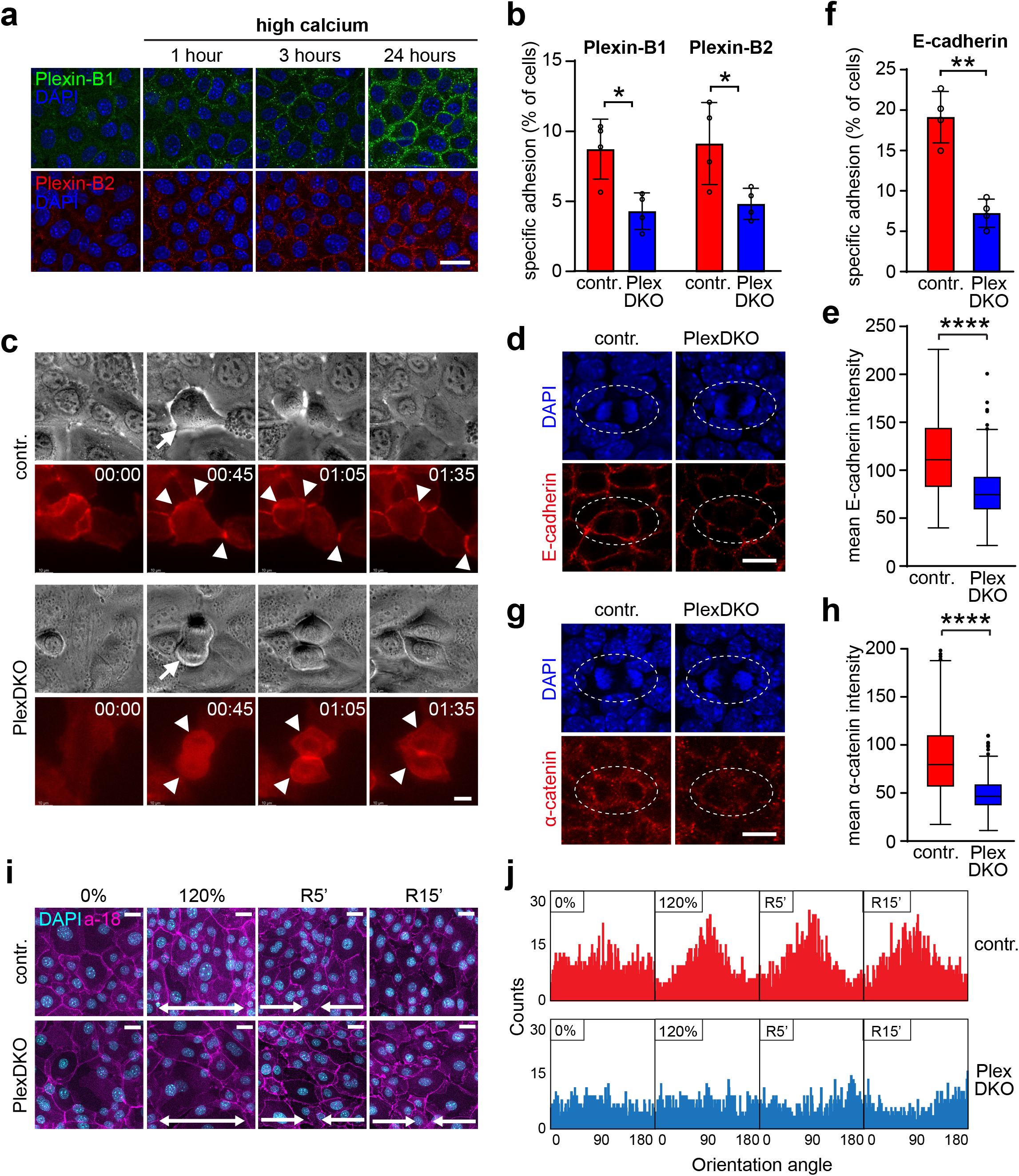
Plexin-B1/Plexin-B2 mediate mechanosensation through stabilization of adhesive cell-cell junctions. **a**, Primary mouse keratinocytes were cultured with 70 μM Ca^2+^. 1.8 mM Ca^2+^ was added (“high calcium”), and cells were analyzed at the indicated time points. Shown are confocal images of immunostainings using anti-Plexin-B1 (green) and Plexin-B2 (red) antibodies. **b**, 96-well nitrocellulose plates were coated with recombinant extracellular portions of murine Plexin-B1 or Plexin-B2, and primary mouse keratinocytes of control or PlexDKO mice were allowed to adhere for 30 min at 37°C. Specific adhesion was quantified as described in Methods. **c**, Live cell imaging of primary mouse keratinocytes engineered to stably express E-cadherin-mRuby. Shown are representative epifluorescence still images of cell divisions. Arrows point to dividing cells; arrowheads point to cell-cell contacts. Time is indicated in an hours:minutes format. Scale bar, 10 μm. **d**, Confocal images of whole-mount immunostainings (top view) of murine epidermis at E15.5 using an anti-E-cadherin antibody (red). Shown are representative images of cell divisions (contoured by dashed lines) in the basal cell layer. **e**, Quantification of E-cadherin immunofluorescence intensities at cell-cell contacts between a dividing cell and its immediate neighbors (box plot with minimum, first quartile, median, third quartile and maximum; control: *n*=207 cell-cell contacts of 34 dividing cells from 3 mice, PlexDKO: *n*=209 cell-cell contacts of 34 dividing cells from 3 mice; Mann-Whitney U test: p<0.0001). **f**, 96-well nitrocellulose plates were coated with the recombinant extracellular portion of murine E-cadherin, and primary mouse keratinocytes of control or PlexDKO mice were allowed to adhere for 30 min at 37°C. Specific adhesion was quantified as described in Methods. **g**, Confocal images of whole-mount immunostainings (top view) of murine epidermis at E15.5 using an anti-α-catenin antibody (red). Shown are representative images of cell divisions in the basal cell layer. **h**, Quantification of α-catenin immunofluorescence intensities at cell-cell contacts between a dividing cell and its immediate neighbors (box plot with minimum, first quartile, median, third quartile and maximum; control: *n*=228 cell-cell contacts of 37 dividing cells from 3 mice, PlexDKO: *n*=244 cell-cell contacts of 39 dividing cells from 3 mice; Mann-Whitney U test: p<0.0001). **i,** Representative α-catenin (a-18 = α-catenin tension-sensitive epitope antibody; magenta) and nuclear (DAPI; cyan) immunofluorescence confocal images of primary mouse keratinocyte monolayers exposed to uniaxial static stretch and subsequent release. Scale bars, 20 μm. **j**, Quantification of immunofluorescence images in (i) showing the orientation angle of cell major axes perpendicular to the stretch direction (frequency distribution of >500 cells/condition pooled across 3 independent experiments; K^2^=366.4 (0% contr.), 5.804 (120% contr.) 164.2 (0% PlexDKO), 215.2 (120% PlexDKO); D’Agostino-Pearson Omnibus test).

Similar to Plexin-B1/Plexin-B2, cadherins are calcium-dependent homophilic adhesion molecules, which act as mechanical integrators upstream of YAP^33–35^. We therefore asked whether plexins could mediate mechanosensation through the regulation of adherens junction organization and/or function. Indeed, we observed that the localization of E-cadherin to cellcell contacts of dividing primary mouse keratinocytes *in vitro* (Fig. 4c) and of dividing epidermal stem cells *in vivo* (Fig. 4d,e) was strongly reduced by Plexin-B1/Plexin-B2-deficiency. Consequently, adhesion of Plexin-B1/Plexin-B2-deficient primary keratinocytes to E-cadherin was impaired (Fig. 4f). E-cadherin forms a complex with α-catenin, which is an upstream negative regulator of YAP and is critically involved in cell density sensing in the epidermis^27, 34, 36^. Fully consistent with the decrease in E-cadherin-based junctions, we found a diminished recruitment of α-catenin to junctions of dividing epidermal stem cells lacking Plexin-B1/Plexin-B2 (Fig. 4g,h).

Finally, to assess whether Plexin-B1/Plexin-B2 mediate mechanosensation through regulation of cell junctions, we analyzed the alignment of keratinocyte monolayers relative to the direction of mechanical stretch, a response known to be junction-dependent^37^. While control cells robustly reoriented their major axes perpendicular to the direction of stretch, Plexin-B1/Plexin-B2 double-deficient cells entirely failed to mount a mechanoresponse (Fig. 4i,j). Collectively, these results indicate that Plexin-B1/Plexin-B2 stabilize adhesive cell junctions required for sensation of mechanical forces.

### Plexin-B1/Plexin-B2 regulate dynamic crowding-induced changes of cortical stiffness

Cadherin-based cell junctions not only connect, via α-catenin, to the cortical actomyosin network, but also contribute to its biogenesis^34^. The cortical actomyosin network, in turn, generates cortical tension and promotes cadherin clustering and stability at cell-cell contact sites^34, 35, 38, 39^. We therefore investigated whether Plexin-B1/Plexin-B2 impact on the organization and regulation of the actomyosin network. Indeed, cortical F-actin intensity at contact sites of dividing epidermal stem cells and their respective neighbor cells was less pronounced in the epidermis of Plexin-B1/Plexin-B2-deficient embryos as compared to control embryos (Fig. 5a,b). Moreover, the levels of phosphorylated myosin light chain 2 (pMLC-2), a marker for actomyosin contractility, were markedly reduced (Fig. 5c,d). In order to probe for a potential requirement of Plexin-B1/Plexin-B2 in the regulation of the actomyosin network by mechanical forces, we exposed control and Plexin-B1/Plexin-B2-knockout keratinocytes to static stretch and abrupt compression. These analyses revealed that, in contrast to control cells, Plexin-B1/Plexin-B2-deficient cells were unable to modulate the levels of pMLC-2 in response to crowding (Fig. 5e,f). We then asked whether this impaired modulation of pMLC-2 levels would correlate with an impaired regulation of cell surface area and cortical stiffness in response to crowding. When released from stretch, control primary mouse keratinocytes rapidly reduced their surface area (Fig. 5g). In striking contrast, keratinocytes lacking Plexin-B1/Plexin-B2 failed to respond by a surface area decrease (Fig. 5g). To examine the crowding-induced regulation of cortical stiffness, we performed atomic force microscopy (AFM)-mediated force indentation spectroscopy on micropatterns at different cell densities. While control cells, as expected, showed a linear inverse correlation of cortical elastic modulus and cell density, Plexin-B1/Plexin-B2-deficient cells exhibited no correlation (Fig. 5h). Taken together, these data demonstrate that Plexin-B1/Plexin-B2 are required for dynamic modulation of the actomyosin network, cell surface area and cortical stiffness in response to lateral compression.

**Fig. 5:**
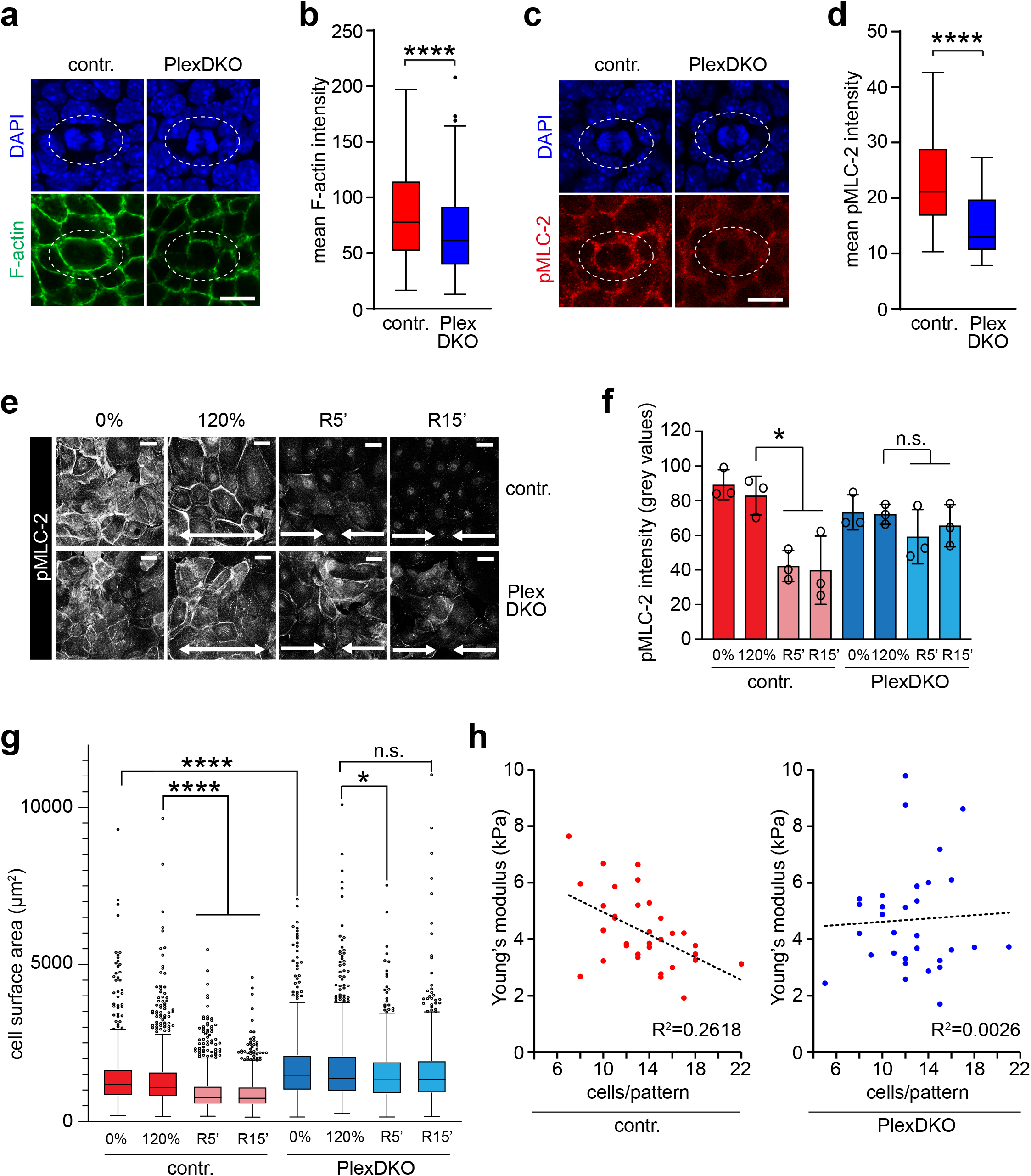
Plexin-B1/Plexin-B2 regulate dynamic crowding-induced changes of cortical stiffness. **a**, Confocal images of whole-mount stainings (top view) of murine epidermis at E15.5 using phalloidin (green). Shown are representative images of cell divisions (contoured by dashed lines) in the basal cell layer. Scale bar, 10 μm. **b**, Quantification of phalloidin fluorescence intensities at cell-cell contacts between a dividing cell and its immediate neighbors (box plot with minimum, first quartile, median, third quartile and maximum; control: *n*=228 cell-cell contacts of 37 dividing cells from 3 mice, PlexDKO: *n*=244 cell-cell contacts of 39 dividing cells from 3 mice; Mann-Whitney U test: p<0.0001). **c**, Confocal images of wholemount immunostainings (top view) of murine epidermis at E15.5 using an anti-phosho-myosin light chain 2 (pMLC-2; Ser19) antibody (red). Shown are representative images of cell divisions (contoured by dashed lines) in the basal cell layer. Scale bar, 10 μm. **d**, Quantification of pMLC-2 immunofluorescence intensities at cell-cell contacts between a dividing cell and its immediate neighbors (box plot with minimum, first quartile, median, third quartile and maximum; control: *n*=50 dividing cells from 3 mice, PlexDKO: *n*=47 dividing cells from 3 mice; Mann-Whitney U test: p<0.0001). **e,f**, Representative pMLC-2 (Ser19) immunofluorescence images (e) and quantification (f) upon crowding (mean **±** s.d., *n*=3 independent experiments with >100 cells/condition/experiment; *p = 0.0322, Friedman/Dunn’s; scale bars 20 μm). **g**, Quantification of cell surface areas after static 120% stretch and subsequent release (Tukey’s box and whiskers plot; n>500 cells/condition pooled across 3 independent experiments; ****p<0.0001, *p=0.0131, Kruskal-Wallis/Dunn’s). **h**, Force indentation spectroscopy of cell cortexes from primary mouse keratinocytes adhering on 100 μm circular micropatterns (control: *n*=34, PlexDKO: *n*=31 micropatterns pooled across 3 independent experiments).

### Plexin-B1/Plexin-B2 control YAP activity and cell proliferation in basal cell carcinoma

Next, we asked whether Plexin-B1/Plexin-B2-mediated mechanosensation of proliferation-induced tissue crowding could also be relevant in skin disease. Consistent with the low rate of stem cell divisions in the adult murine epidermis (Fig. 1a and^19, 20^), genetic inactivation of Plexin-B1 and Plexin-B2 at adult stages using a tamoxifen-inducible K14-CreERT line (genotype K14-CreERT;*plxnb1*^flox/flox^;*plxnb2*^flox/flox^; Supplementary Fig. 5a) did not result in any morphological abnormalities (Supplementary Fig. 5b,c). We hypothesized that an experimental triggering of cell divisions in the adult by topic application of a chemical mitogen to the skin would again disclose the requirement of Plexin-B1/Plexin-B2 for the inhibitory mechanical feedback on cell proliferation. Indeed, treatment of Plexin-B1/Plexin-B2-deficient mice with the chemical mitogen, the phorbol ester TPA (Supplementary Fig. 5d) resulted in higher proliferation rates (Supplementary Fig. 5e,f) and more pronounced epidermal thickening (Supplementary Fig. 5g,h) than in control mice. Fully in line with the function of Plexin-B1/Plexin-B2 as mechanosensitive suppressors of YAP activity during skin development, YAP was also found to be disinhibited in the TPA-treated skin of adult Plexin-B1/Plexin-B2 knockout mice (Supplementary Fig. 5i,j). To assess the potential pathophysiological relevance of mechanosensation through Plexin-B1/Plexin-B2 in disease, we next analyzed the role of Plexin-B1/Plexin-B2 in skin tumors. To do so, we employed a genetic mouse model, in which tamoxifen-inducible inactivation of the tumor suppressor Gαs in the epidermis promotes stem cell proliferation and formation of basal cell carcinomas, the most common type of skin cancer^40^ (Fig. 6a). Interestingly, mice lacking Plexin-B1 and Plexin-B2 displayed a higher incidence (Fig. 6b) and larger sizes of tumors (Fig. 6c,d) than the respective control mice. Furthermore, disruption of Plexin-B1 and Plexin-B2 resulted in a strong increase in cell proliferation (Fig. 6e,f) and a marked up-regulation of YAP activity (Fig. 6g,h). In summary, these data indicate that Plexin-B1/Plexin-B2 limit YAP activity and proliferative capacity not only during embryonic development but also in skin cancer.

**Fig. 6:**
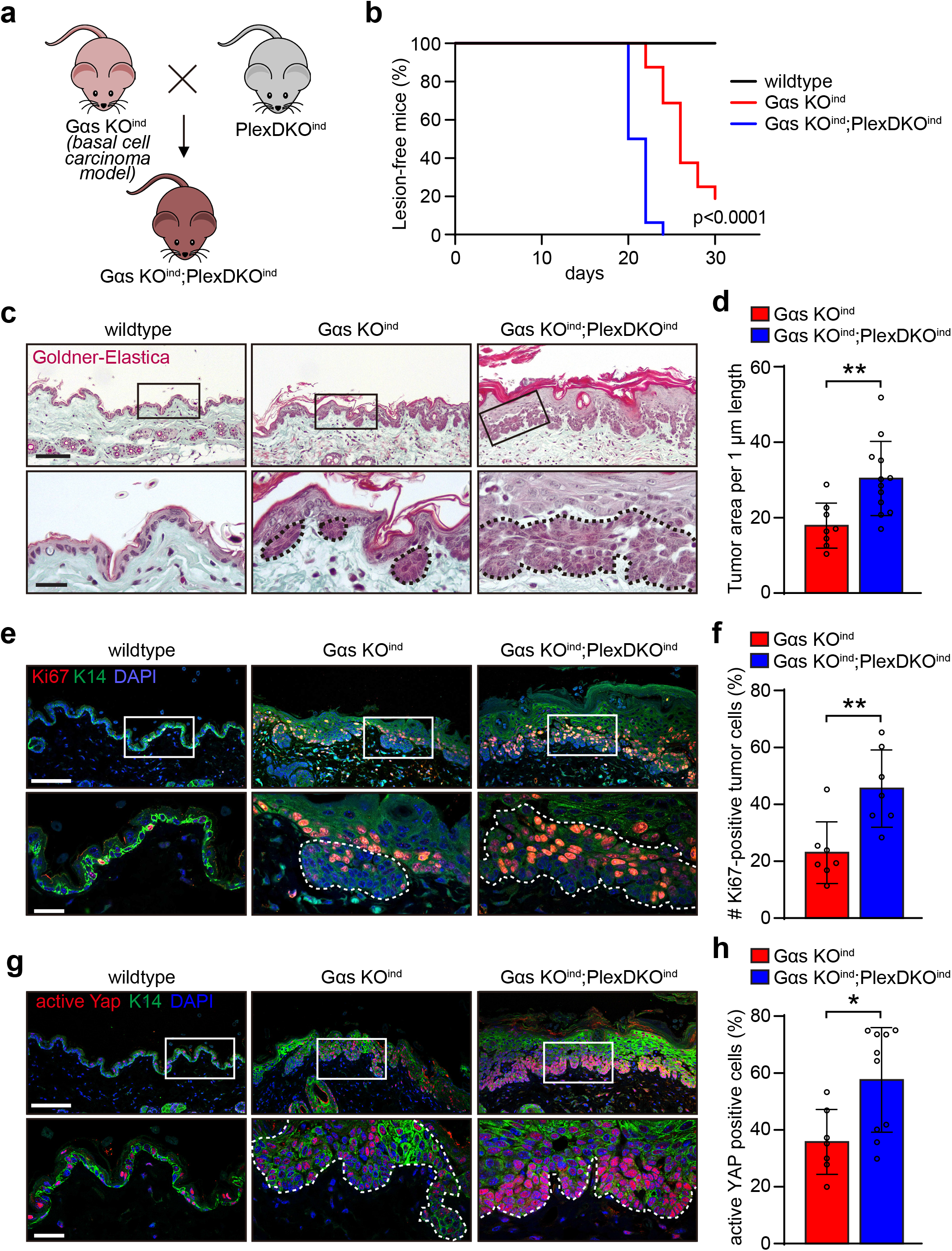
Plexin-B1/Plexin-B2 control YAP activity and cell proliferation in basal cell carcinoma. **a**, Schematic illustration of the generation of epidermis-specific tamoxifen-inducible Gαs/Plexin-B1/Plexin-B2 triple-deficient mice. **b**, Kaplan-Meier curves representing the percentage of lesion-free mice. Time point “0” indicates the start of tamoxifen treatment (wildtype mice: *n*=3, Gαs KO^ind^ mice: *n*=16, Gαs KO^ind^;PlexDKO^ind^ mice: *n*=16; Mantel-Cox test for Gαs KO^ind^ compared to Gαs KO^ind^;PlexDKO^ind^). **c-h**, Histological/immunofluorescence stainings (left panels) and respective quantifications (right panels) of adult murine skin of mice with the indicated genotypes 30 days after treatment with tamoxifen. Boxed areas are magnified in the lower rows. Tumors are marked by dashed lines. **c**, Goldner-Elastica stain of histological sections. Scale bar, 100 μm (upper row), 25 μm (lower row). **d**, Quantification of tumor areas (mean ± s.d.; Gαs KO^ind^ mice: *n*=8, Gαs KO^ind^;PlexDKO^ind^ mice: *n*=12; unpaired t-test). **e**, Confocal images of immunostainings using anti-Ki67 (red) and anti-K14 antibodies (green). Scale bar, 100 μm (upper row), 25 μm (lower row). **f**, Quantification of Ki67-positive cells (mean ± s.d.; Gαs KO^ind^ mice: *n*=7, Gαs KO^ind^;PlexDKO^ind^ mice: *n*=7; unpaired t-test). **g**, Confocal images of immunostainings using anti-active YAP (red) and anti-K14 antibodies (green). Scale bar, 100 μm (upper row), 25 μm (lower row). **h**, Quantification of cells positive for active YAP (mean ± s.d.; Gαs KO^ind^ mice: *n*=7, Gαs KO^ind^;PlexDKO^ind^ mice: *n*=10; unpaired t-test).

## DISCUSSION

Cell divisions within the jammed basal layer of the mouse embryonic epidermis result in crowding and mechanical stress anisotropy. Our study is in line with a concept, in which crowding acts as a central checkpoint in the control of epidermal morphogenesis by triggering mechanisms to reinstate stem cell density. These mechanisms include differentiation and extrusion of differentiating cells from the stem cell layer by upward efflux^18, 41–43^ as well as suppression of stem cell divisions by mechanical feedback^36^. Our data now identify the transmembrane receptors Plexin-B1 and Plexin-B2 as key to this feedback by coupling mechanosensation of cell density to the biochemical control of YAP activity and cell proliferation. Our work demonstrates that the Plexin-B1/Plexin-B2-mediated mechanoresponse to crowding is particularly relevant for proper morphogenesis of the early embryonic epidermis where proliferation rates are high. The sharp decrease of Plexin-B1 and Plexin-B2 expression levels at late embryonic stages and, in particular, the absence of epidermal abnormalities after induction of Plexin-B1/Plexin-B2 deficiency in adult mice indicate that – in contrast to early development – the mechanosensation of crowding does not coordinate epidermal homeostasis when proliferation rates are low. This is consistent with previous studies showing that in the adult interfollicular epidermis, stem cells divide only upon demand to replace delaminating cells, and thus crowding does not occur^10, 44^. Intriguingly, we find that mechanosensation through Plexin-B1/Plexin-B2 regains biological relevance in the adult epidermis for the suppression of cancer cell proliferation. This strongly suggests that loss of Plexin-B1/Plexin-B2 could represent a mechanistic basis for the loss of contact inhibition, a hallmark of cancer that has remained poorly understood on a molecular level^45, 46^.

The extracellular domain of several plexins has been shown to adopt different conformations ranging from a nearly closed, ring-like form to a more-open, chair-like form^17, 47, 48^. Very recently, a particular plexin, Plexin-D1, has been reported to sense shear stress-induced mechanical forces in endothelial cells^17^. This function in mechanosensation required the opening of the ring-like conformation of the Plexin-D1 extracellular domain, but was independent of the binding of semaphorins^17^. This is in line with our findings in epithelial cells, showing that activation of Plexin-B1 and Plexin-B2 by semaphorins is not sufficient to regulate the activity of the mechanotransducer Yap. The similar structures of the extracellular domains of plexins^13, 47^ suggest that the mechanism of mechanosensation by semaphorin-independent conformational changes could be shared by different plexins including Plexin-B1 and Plexin-B2.

In summary, our work uncovers a critical requirement for Plexin-B1 and Plexin-B2 in the control of epidermal stem cell proliferation by mechanical forces. Whether the function of Plexin-B1 and Plexin-B2 in mechanosensation is specific for crowding-induced compression remains open to future research; however, it seems more likely that they could respond to a wider spectrum of different mechanical stimuli, a property that is known from other types of mechanosensors such as mechanosensitive ion channels^49^. Given the wide expression of Plexin-B1 and Plexin-B2 in epithelial and non-epithelial cells both during development as well as in the adult^13, 50, 51^, it is tempting to speculate that mechanosensation by Plexin-B1 and Plexin-B2 could be instrumental in physiology and pathophysiology of multiple tissues.

## ACKNOWLEDGEMENTS

We thank Fatma Aktuna, Eva Braun, Marga Losekam, Andrea Wüstenhagen and Martina Finkbeiner for technical assistance. We thank Dr. Merkel and his team, Institute of Biological Information Processing, Mechanobiology (IBI-2), Forschungszentrum Jülich, Jülich, Germany, for providing the uniaxial stretching device. This work was supported by a grant of the Deutsche Forschungsgemeinschaft to T.W. (GRK 2213), by grants of the Deutsche Forschungsgemeinschaft (73111208 - SFB 829) and the Juselius Foundation to S.A.W., and by a grant of the China Scholarship Council (CSC) to L.Z. (201908080171).

## AUTHOR CONTRIBUTIONS

C.J., A.J., L.K., M.M.N., D.Z., D.T.B., J.F., L.Z., and C.H. performed experiments and analysed data. K.S., A.O., and D.M. provided anti-Plexin-B1 antibodies and recombinant Plexin-B1. L.S.W., C.M.N. and S.O. provided genetically-modified mouse lines. C.M.N. helped with establishing primary mouse keratinocyte cultures. C.J., S.W. and T.W. designed experiments and interpreted data. T.W. conceived and supervised the study and wrote the paper. All authors read and corrected the manuscript.

## METHODS

### Mice

To generate mice lacking Plexin-B1 and Plexin-B2 specifically in the epidermis, mice carrying conditional alleles of Plexin-B1^52^ and Plexin-B2^52^ were crossed with mice expressing Cre constitutively under the control of the keratin 14 (K14) promoter^24^, or with mice expressing CreERT under the control of the K14 promoter^53^ (Tg(KRT14-cre/ERT)20Efu/J, The Jackson Laboratory, Stock No. 005107). Mice carrying floxed alleles of Gαs have been described previously^54^. All mice used in this study were on a C57BL/6 genetic background. Mice were housed under a 12-h light-dark cycle with free access to food and water, and under specific pathogen-free conditions. All procedures were performed in accordance with German Animal Welfare legislation.

### Antibodies

The following primary antibodies were used: guinea pig polyclonal anti-keratin 14 (1:200, Progen, cat. no. GP-CK14), rabbit polyclonal anti-keratin 14 (1:200, Biolegend, cat. no. #905301), rabbit polyclonal anti-keratin 10 (1:200, Biolegend, cat. no. #905401), rabbit polyclonal anti-Ki67 (1:200, Abcam, cat. no. ab15580), rabbit polyclonal anti-phospho-histone H3 (1:200, Cell Signaling, cat. no. #9701), armenian hamster monoclonal anti-Plexin-B2 (for immunostainings; 1:200, eBioscience, cat. #14-5665-85), sheep polyclonal anti-Plexin-B2 (for Western Blot; 1:500, R&D Systems, cat. no. AF6836), rabbit monoclonal anti-E-cadherin (1:200, Cell Signaling, cat. no. #3195), rabbit polyclonal anti-α-catenin (1:200, Invitrogen, cat. no. 71-1200), anti-α-catenin a-18 (for immunostainings on elastomers; 1:10000,^55^), rabbit monoclonal anti-cleaved Notch1 (NICD, 1:100, Cell Signaling, cat. no. #4147), rabbit monoclonal anti-Yap antibody (1:200, Cell Signaling, cat. no. #14074), mouse monoclonal anti-Yap (for immunostainings on elastomers; 1:300, Santa Cruz; sc-101199), rabbit monoclonal anti-active Yap (1:200, Abcam, cat. no. ab205270), rabbit polyclonal anti-phospho-myosin light chain 2 (Ser19) (1:200, Cell Signaling, cat. no. #3671), rabbit polyclonal anti-phospho-myosin light chain 2 (Thr18/Ser19) (for immunostainings on elastomers; 1:200, Cell Signaling; cat. no. #3674). An anti-Plexin-B1 antibody was raised against the extracellular domain of Plexin-B1. The following secondary antibodies were used: Cy3-conjugated anti-armenian hamster (1:200; Jackson ImmunoResearch, cat. no. 127-165-160), AlexaFluor 488-conjugated anti-guinea pig (1:200; Invitrogen, cat. no. A11073), AlexaFluor 488-conjugated anti-rabbit (1:200; Invitrogen, cat. no. R37118), AlexaFluor 555-conjugated anti-rabbit (1:200; Invitrogen, cat. no. A21429), AlexaFluor 555-conjugated anti-mouse (1:200; Invitrogen, cat. no. A21424), AlexaFluor 488-conjugated anti-mouse (1:200; Invitrogen, cat. no. A11029), HRP-conjugated anti-sheep (1:5000, Invitrogen, cat. no. A16041).

The following reagents were used: AlexaFluor 488-conjugated phalloidin (1:500, Invitrogen, cat. no. A12379), AlexaFluor 647-conjugated phalloidin (1:500, Invitrogen, cat. no. A22287).

### Recombinant proteins

Recombinant mouse Sema4C (cat. no. 8394-S4-050), recombinant mouse Plexin-B2 (cat. no. 6836-PB-050), and recombinant mouse E-cadherin (cat. no. 8875-EC-050) were purchased from R&D Systems. A fragment of human Plexin-B1 cDNA (encoding amino acids 1-535) was cloned into the pcDNA5 vector using HindIII (R0104S, NEB) and XhoI (R0146S, NEB) restriction sites with addition of a C-terminal 6xHis tag (KHHHHHH). The expression plasmid was transfected into Expi293F cells (Invitrogen) according to the manufacturer’s instructions. The supernatant containing secreted Plexin-B1 (amino acids 20-535)-6xHis was collected after 7 days and purified using a 2-step purification protocol developed at Medical Research Council Technology (MRC-T). Briefly, overexpressed protein was captured on an Excel HisTrap (GE) column in 20 mM HEPES, pH 8.0, 0.3 M NaCl and 10 mM imidazole buffer. After washing the column with 10 column volumes of washing buffer (20 mM HEPES, pH 8.0, 0.3 M NaCl and 20 mM imidazole), the protein was eluted with elution buffer (20 mM HEPES, pH 8.0, 0.3 M NaCl and 250 mM imidazole), followed by gel filtration chromatography using a Superdex 200 16/60 column equilibrated with PBS. Collected fractions were analysed on SDS-PAGE, pooled, quantified by NanoDrop, aliquoted and flash-frozen for storage at −80°C.

### Quantitative RT-PCR

Total RNA from murine skin, murine epidermis, and primary mouse keratinocytes was isolated using Direct-zol RNA Microprep Kits (Zymo) according to the manufacturer’s instructions. Genomic DNA was removed by on-column DNAse digestion, and cDNA was generated by reverse transcription using the RevertAid First Strand cDNA Synthesis Kit (ThermoFisher). qRT-PCR was performed with SYBR Green Supermix (BioRad) in a Real Time Quantitative Thermal Cycler (BioRad). For absolute quantifications, mouse genomic DNA was used as a standard. For relative quantifications, gene expression changes were normalized to *gapdh* expression.

The following primers were used for absolute quantification of gene expression: for murine *plxna1*: forward 5’-GGGATGCTGCAGGTGTATTC-3’ and reverse 5’-CACCACCGATACCCACAAT-3’; for murine *plxna2*: forward 5’-GGACTATGAGCTCCACAGTGATT-3’ and reverse 5’-AGGGTGTCAGAGGGAATCTTG-3’; for murine *plxna3*: forward 5’-GTTACAGCTGCTGGTCATGC-3’ and reverse 5’-GCACCCTCCTATGGTGAAGA-3’; for murine *plxna4*: forward 5’-CAGCATGCAGGCTTTGTG-3’ and reverse 5’-AAGAGCGGGCCATGAACT-3’; for murine *plxnb1*: forward 5’-CTGCCAGGGTGGTCGTTA-3’ and reverse 5’-ACCTCCTTGGACGTGGCTA-3’; for murine *plxnb2*: forward 5’-CAAACACCAGGTGGAAAAGG-3’ and reverse 5’-GCCTGTGTCATTGAGGGTGT-3’; for murine *plxnb3*: forward 5’-GCCAGGCCTTTAATGATGTG-3’ and reverse 5’-AGAAGGGTCAGGGTTGCAG-3’; for murine *plxnc1*: forward 5’-GCCAGATTCAGAAGCATTCC-3’ and reverse 5’-TGAACTT-GTGTTTCCCTCGAT-3’; for murine *plxnd1*: forward 5’-CGGTTCCGCCTAGACTACCT-3’ and reverse 5’-TGGGTGATGTTTGATCCACTT-3’. The following primers were used for relative quantification of gene expression: for murine *ctgf*: forward 5’-TGACCTGGAGGAAAACATTAAGA-3’ and reverse 5’-AGCCCTGTATGTCTTCACACTG-3’; for murine *cyr61*: forward 5’-GGATCTGTGAAGTGCGTCCT-3’ and reverse 5’-CTGCATTTCTTGCCCTTTTT-3’; for murine *gapdh*: forward 5’-AGGTCGGTGTGAACGGATTTG-3’ and reverse 5’-TGTAGACCATGTAGTTGAGGTCA-3’.

### Cell lines

MDCK cells, a canine renal tubular epithelial cell line, were cultured in MEM supplemented with 10% FBS^23^.

### Primary mouse keratinocytes

Primary mouse keratinocytes were isolated and cultured essentially as described^56^. Briefly, the skin of newborn mice was harvested using sterile forceps, and incubated in 1 ml dispase (5mg/ml) in culture medium (CnT-07; Cellntec) in a 2-ml-microcentrifuge tube over night at 4°C. The skin was then transferred to a 10-cm-petri dish, and washed with PBS. Using forceps, the epidermal sheet was separated from the dermis, transferred with the basal layer facing downwards onto 500 μl of accutase (Cellntec) in a 6-cm-petri dish, and incubated for 30 min at RT. Keratinocytes were washed out of the epidermal basal layer using 2 ml of culture medium, collected by centrifugation, and seeded onto collagen type 1 (30ug/ml, Biochrom, cat no. L7213)-coated 10-cm-petri dishes. Primary mouse keratinocytes were kept in culture medium (CnT-07; Cellntec) at 37°C and 5% CO2. Only low passage cells (max. 2 passages) were used for experiments. For the analysis of semaphorin-plexin forward and reverse signaling, primary mouse keratinocytes were incubated with recombinant proteins (25 nM) for 8 hours prior to RNA isolation.

### Lentiviral and retroviral transduction

To generate an E-cadherin-mRuby lentiviral construct, the human cDNA of E-cadherin was amplified by PCR and fused to the mRuby2 sequence using overlap/extension PCR. The fused cDNA was ligated into pWPXL via Mlu1 and Spe1 restriction sites. A pWPXL lentiviral construct encoding pEGFP-α-catenin was a kind gift from Katharina Grikscheit^57^. For lentivirus production, HEK293T cells were cotransfected (using the calcium phosphate method) with the packaging plasmids psPAX2 and pMD2.G together with the aforementioned lentiviral constructs. After 48 hours, supernatants containing viral particles were harvested, filtered, and used to transduce primary mouse keratinocytes.

To generate MDCK cells stably expressing Plexin-B2-EGFP or expressing a Plexin-B2 mutant, which lacks the intracellular domain (Plexin-B2ΔIC-EGFP), the murine cDNAs encoding triple-myc-tagged wildtype Plexin-B2 (amino acids 20 – 1842)^58^ or triple-myc-tagged truncated Plexin-B2 (amino acids 20 – 1227) were cloned into pEGFP-N1 using AfeI and SalI restriction sites. The fused cDNAs encoding Plexin-B2 and EGFP were ligated into pLNCX2 using NotI. For retrovirus production, PT67 cells were transfected with the aforementioned retroviral constructs. After 48 hours, supernatants containing viral particles were harvested, filtered, and used to transduce MDCK cells (in the presence of polybrene; 8 μg/ml).

### H&E and Goldner-Elastica staining

H&E and Goldner-Elastica stainings were performed on paraffin-embedded sections using standard laboratory protocols.

### Immunofluorescence staining

For whole-mount immunofluorescence stainings, the epidermis was harvested from the dorsal skin of E15.5 embryos using forceps, fixed in 4% PFA for 30 minutes, washed twice in 50 mM glycine in PBS, followed by washing twice in PBS. After permeabilization with PBST (0.5% Triton X-100 in PBS) for 20 minutes and blocking with 4% FCS in PBST for at least 1 hour, the epidermis was incubated with primary antibodies (diluted in 2% FCS in PBST) at 4°C overnight. After three times washing in PBST, samples were incubated with secondary antibodies and DAPI for 3 hours, and washed. All processing of samples was done in 12- or 96-well-plates. Samples were then mounted on glass slides using Fluoromount (Dako). Images were taken using a confocal laser scanning microscope (Zeiss LSM 700).

For immunofluorescence stainings of paraffin-embedded sections, tissues (dorsal skin of mouse embryos or adult mouse skin/tumor tissue) were fixed in 4% PFA at 4°C overnight, embedded in paraffin according to standard laboratory protocols, sectioned at 5 μm, and mounted on glass slides. Antigen retrieval was done by boiling in a steam pressure pot in 10 mM sodium citrate buffer pH 6.0 for 10 minutes. Sections were permeabilized with PBST (0.2% Triton X-100 in PBS) for 10 minutes, blocked with 4% FCS in PBST for at least 1 hour and incubated with primary antibodies (diluted in 2% FCS in PBST) at 4°C overnight, washed with PBS, and incubated with secondary antibodies and DAPI at room temperature for 1 hour. After three times washing in PBS, sections were mounted with Fluoromount (Dako). Images were taken using a confocal laser scanning microscope (Zeiss LSM 700).

For immunofluorescence staining of cryotome sections, dorsal skin of mouse embryos or adult mouse skin were fixed in 0.2% PFA overnight at 4°C, followed by one day of incubation in 30% sucrose (in PBS). Tissues were embedded in O.C.T. medium, frozen on dry ice, and sectioned using a cryotome (thickness of sections 25 μm), and mounted on glass slides. Sections were postfixed in 4% PFA at 4°C for 30 minutes, washed twice in 50 mM glycine in PBS, permeabilized with PBST (0.2% Triton X-100 in PBS), blocked with 4% FCS in PBST for at least 1 hour and incubated with primary antibodies (diluted in 2% FCS in PBST) overnight at 4°C. Sections were washed three times in PBS, incubated with secondary antibodies and DAPI at room temperature for 1 hour. After three times washing in PBS, sections were mounted with Fluoromount (Dako). Images were taken using a confocal laser scanning microscope (Zeiss LSM 700).

For immunofluorescence stainings of primay mouse keratinocytes, keratinocytes (seeded into μ-Slide 8 well (ibidi) or onto micropatterns) were fixed in 4% PFA for 30 minutes, washed 3 times with PBS, permeabilized with PBST (0.2% Triton X-100 in PBS) for 10 minutes, blocked with 4% FCS in PBST for at least 1 hour, incubated with primary antibodies (diluted in 2% FCS in PBST) at 4°C overnight, washed with PBS and incubated with secondary antibodies and DAPI at room temperature for 1 hour. After three times washing in PBS, cells were mounted with Fluoromount (Dako). Images were taken using a confocal laser scanning microscope (Zeiss LSM 700).

For immunostainings on silicone elastomers, cells were fixed in 4% PFA for 10 min room temperature, permeabilized with 0.3% Triton X-100 in PBS, and blocked in 5% bovine serum albumin (BSA). Samples were subsequently incubated overnight in primary antibody in 1% BSA/0.3% Triton X-100/PBS, followed by washing in PBS and incubation in secondary antibody in 1% BSA/0.3% Triton X-100/PBS. Finally, samples were mounted in Elvanol. Images were collected by laser scanning confocal microscopy (SP8X; Leica) with Leica Application Suite software (LAS X version 2.0.0.14332), using 40x immersion objectives. Images were acquired at room temperature using sequential scanning of frames of 1 μm thick confocal planes (pinhole 1) after which 10 planes encompassing complete cell nuclei were projected as a maximum intensity confocal stack. Images were collected with the same settings for all samples within an experiment.

### Immunohistochemical staining

For immunohistochemical stainings of paraffin-embedded sections, tissues were fixed in 4% PFA at 4°C overnight, embedded in paraffin according to standard laboratory protocols, sectioned at 5 μm, and mounted on glass slides. Antigen retrieval was done by boiling in a steam pressure pot in 10 mM sodium citrate buffer pH 6.0 for 10 minutes. For immunohistochemical staining of NICD, sections were incubated with BLOXALL (ImmPRESS Excel Staining Kit; Vector laboratories, cat. no. MP-7601) to inactivate endogenous peroxidases for 10 minutes, washed in PBS, permeabilized with PBST (0.2% Triton X-100 in PBS) for 10 minutes, blocked with 4% FCS in PBST for at least 1 hour, and incubated with anti-NICD antibody (diluted in 2% FCS in PBST) at 4°C overnight, washed with PBS, followed by usage of the ImmPRESS Excel Staining Kit (Vector laboratories, cat. no. MP-7601). Sections were then dehydrated and mounted with Pertex.

### TUNEL assay

TUNEL assays were performed on paraffin-embedded sections of the dorsal skin of embryos using the DeadEnd Fluorometric TUNEL System (Promega, cat. no. G3250) according to the manufacturer’s instructions.

### Surface biotinylation assay

Confluent primary mouse keratinocytes in 10-cm petri dishes were washed twice with ice-cold PBS followed by incubation with 8 ml of Biotin-NHS (Sigma, cat. no. 203112) in PBS (0.25 mg/ml) for 10 min at 4 °C with gentle shaking. Afterwards, the reaction was quenched with the addition of 6 ml Tris-HCl (50 mM, pH 7,4). Cells were scraped off the dish, collected in a 50 ml tube, pelleted by centrifugation at 4 °C, and washed 3 times with ice-cold PBS. The pellet was resuspended in 800 μl of RIPA buffer (50 mM Tris-HCl, 300 mM NaCl, 1 mM EDTA, 1% Triton X-100, 0.25% sodium deoxycholate; 0.1% SDS, 5 mM MgCl_2_; pH 7.4) containing cOmplete Protease Inhibitor Cocktail (Roche, cat. no. 4693132001), and sonicated three times. After centrifugation at 4 °C, supernatants were incubated with Pierce Streptavidin Agarose (ThermoFisher, cat. no. 20353) for 90 min at 4 °C. Agarose beads were washed 4 times with ice-cold lysis buffer, and boiled in Laemmli buffer before analysis on SDS-PAGE followed by a Western blotting.

### Fluorescence in situ hybridization (FISH)

Fluorescence in situ hybridizations (FISH) were done on paraffin-embedded sections (5 μm) using the RNAscope system (ACD). The following probes were used: Mm-CTGF (ACD, cat. no. 314541), Positive Control Mm-Ppib (ACD, cat. no. 313911), Negative Control DapB (ACD, cat.no. 310043). The following kit was used: RNAscope 2.5 HD Detection Reagents-RED (ACD, cat. no. 322360). FISH was combined with immunofluorescence staining as described above.

### Live cell imaging

Live cell imaging of primary mouse keratinocytes was performed using a Thunder Imaging System (Leica) at 37°C in a CO2 humidified incubation chamber in cell culture medium (CnT-07, Cellntec).

### Adhesion assay

Adhesion assays were performed essentially as described^32^. Mixed Cellulose Ester 96-well plates (Millipore, cat. no. MSHAS4510) were coated without or with recombinant proteins (5 μg/ml) at 4 °C overnight. Wells were then blocked with 5% bovine serum albumin (BSA) for 1 hour at room temperature. After washing with PBS, 1×10^4^ primary mouse kerationocytes (in culture medium with 1.8 mM calcium) were added per well, and allowed to adhere for 30 minutes at 37°C and 5% CO2. Afterwards, non-adherent cells were removed by washing twice with PBS. For some control wells, non-adherent cells were not removed by washing and all seeded cells remained in the well. The CellTiter-Glo Luminescent Cell Viability Assay (Promega, cat. no. G7572) was used according to the manufacturer’s protocol, and the percentage of specific adhesion was quantified using the following formula: specific adhesion [%] = RLU of coated well after removal of non adherent cells – RLU of non coated well after removal of non adherent cells

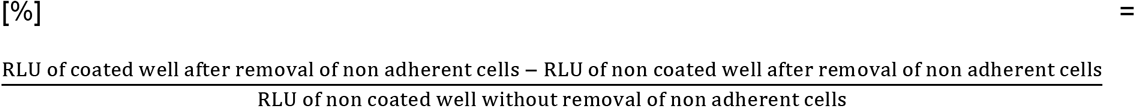

### Pharmacological induction of epidermal proliferation

Mice at the age 6 weeks were injected with tamoxifen (1 mg; i.p.) for 5 consecutive days. One week after the start of tamoxifen injections, the dorsal hair was shaved. Two days later, DMBA (25 μg = 97.5 nmol, dissolved in acetone) was applied to the shaved back skin. Starting two weeks after the treatment with DMBA, 12-O-Tetradecanoylphorbol-13-acetate (TPA; 4 μg = 6.5 nmol, dissolved in acetone) was applied to the same site twice per week for a total of 14 weeks. The analyzed genotypes were “contr.” (genotype *plxnb1*^flox/flox^;*plxnb2*^flox/flox^) and “PlexDKO^ind^” (genotype K14-CreERT;*plxnb1*^flox/flox^;*plxnb2*^flox/flox^). Mice were regularly monitored for the appearance of skin lesions, and did not develop papillomas. Only female mice were examined.

### Basal cell carcinoma mouse model

Mice at the age of 7.5 weeks were injected with tamoxifen (1 mg; i.p.) for 5 consecutive days. The analyzed genotypes were “Gαs KO^ind^” (genotype K14-CreERT;*gnas*^flox/flox^;*plxnb1*^+/flox^;*plxnb2*^+/flox^) and “Gαs KO^ind^;PlexDKO^ind^” (genotype K14-CreERT;*gnas*^flox/flox^;*plxnb1*^flox/flox^;*plxnb2*^flox/flox^). Mice were regularly monitored for the appearance of skin lesions. The total observation period (from the start of tamoxifen injections to analysis) was 30 days. Both male and female mice were examined.

### Quantitative image analysis

Segmentation analyses were done based on E-cadherin immunofluorescence stainings using the Tissue Analyzer plug-in in ImageJ. Cell area and shape (major axis/minor axis; roundness: 4x[area] / (π x [major axis]^2^) were quantified by using the “area” and “shape description” functions of ImageJ. The number of cells per area (cell density) was quantified manually.

To quantify mean fluorescence intensities, the segmented line tool of Fiji (line width 5 pixels) was used.

All quantifications were done by an observer blinded to genotypes.

For crowding experiments, images were analyzed using Fiji^59^. To quantify nuclear YAP, fields were randomly selected based exclusively on the presence of nuclei, as assessed by DAPI staining. Areas of interest were generated using automated thresholding of the DAPI staining, after which mean fluorescence intensities of nuclear YAP intensity was quantified within the areas of interest from maximum projection images obtained as described above. The measured mean nuclear intensities were normalized to DAPI intensity of the corresponding individual nuclei to account for possible unevenness in sample topology. To measure pMLC-2 intensity, pMLC-2 signal intensity at junctions was measured by automatic thresholding followed by subtraction of cytoplasmic background signal. Cell area was measured by manually tracing α18 fluorescence. Cell orientation was quantified by measuring the orientation angle (the angle between the long axis and a line parallel to the x-axis of the image which was aligned to the direction of stretch) of the major Feret diameter of areas of interest corresponding to traced α-18 positive adherens junctions.

### Micropatterns

Micropatterned adhesive surfaces were generated using the PRIMO optical module (Alvéole, France) controlled by the Leonardo plugin (V3.3, Alveole) mounted on a Nikon TI-E inverted microscope (Nikon Instruments) equipped with a Super Plan FLuor 20x ELWD lens (Nikon) lens and a DMD-based UV (375nm). To generate circular and square micropatterns, both 100 μm diameter, were projected onto plasma-cleaned (Corona Treater, ETP), PLLgPEG-passivated (0.1 mg/ml PLL-g-PEG (PLL (20)-g [3.5]-PEG (2), SuSoS) 35mm glass-bottom dishes (Ibidi). Patterned areas were then washed multiple times with PBS and conjugated with a uniform coating of 10 μg/ml fibronectin and 38.75 μg/ml collagen for 1h at +37 °C. The substrates were then washed with PBS, prior to seeding 100 K of mouse keratinocytes onto each 35mm dish. Keratinocyte monolayers were allowed to proliferate on the patterns for 24 hours, at which time they were subjected to atomic force microscopy or fixed and processed for immunofluorescence and quantification analyses. For the analysis of YAP localization (Fig. 3d,e), only micropatterns with similar cell densities were compared.

### Crowding experiments

The custom-built uniaxial cell stretcher has been described in detail previously^37, 60^. Polydimethylsiloxane (PDMS) elastomers were prepared from a two-component formulation (Sylgard 184, Dow Corning) by mixing base and crosslinker in a ratio of 40 to 1 (weight/weight) to obtain substrates with 50 kPa stiffness. Chambers were UV sterilized for 1h and finally coated with fibronectin (20 μg/ml) in phosphate-buffered saline (PBS) for 1 h at 37°C prior to cell seeding. 350 000 cells per elastomer (4 cm^2^) were seeded 16 h before experiment start. Before initiation of experiments, culture medium was replaced by medium containing 200 μM Ca^2+^ to promote cell-cell contact formation. Cells were then exposed to 20 % static stretch for 12h and fixed with 4% PFA either directly when under stretch or 5 or 15 min post release.

### Atomic force microscopy (AFM)

AFM measurements were performed on cells cultured on plastic dishes as monolayers or on micropatterns generated on glass bottom dishes (35 mm Ibidi). Measurements were done using a JPK NanoWizard 2 (Bruker Nano) atomic force microscope mounted on an Olympus IX73 inverted fluorescent microscope (Olympus) and operated via JPK SPM Control Software v.5. Spherical silicon dioxide beads with a diameter of 3.5 μm glued onto tipless silicon nitride cantilevers (NanoAndMore, CP-PNPL-SiO-B-5) with a nominal spring constant of 0.08 N m-1 were used. For all indentation experiments, forces of up to 4 nN were applied, and the velocities of cantilever approach and retraction were kept constant at 2μm s–1 ensuring detection of elastic properties only. All analyses (>50 cells per experiment/condition) were performed with JPK Data Processing Software (Bruker Nano). Prior to fitting the Hertz model to obtain cell Young’s Modulus (Poisson’s ratio of 0.5), the offset was removed from the baseline, contact point was identified, and cantilever bending was subtracted from all force curves.

### Statistics

*p ≤ 0.05, **p ≤ 0.01 for unpaired t-tests.

## SUPPLEMENTARY FIGURE LEGENDS

**Supplementary Fig. 1:**
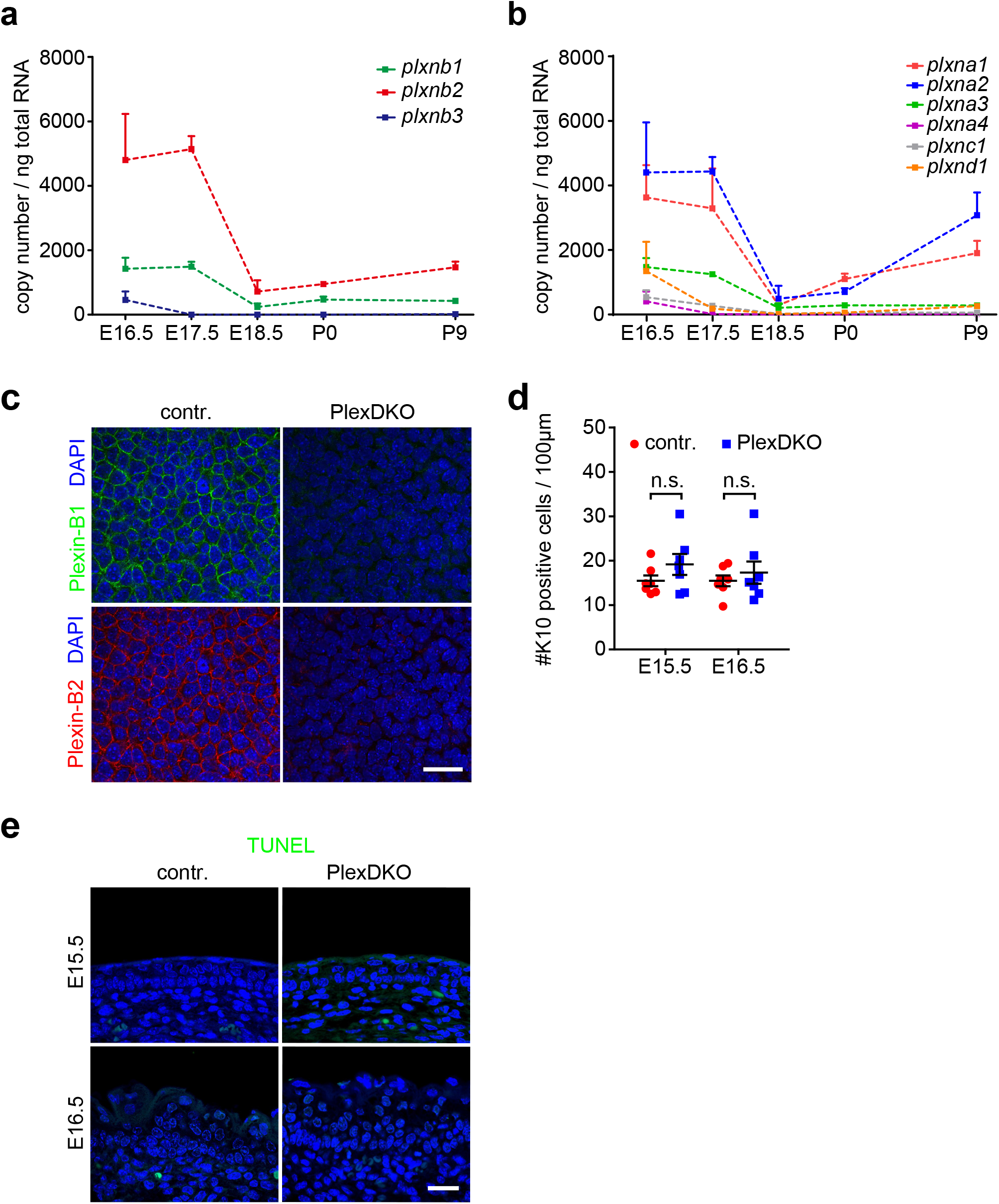
**a,b**, mRNA expression levels in the murine epidermis of genes encoding plexins (*plxn*) at different time points of embryonic (E) and postnatal (P) development as determined by quantitative RT-PCR (mean ± s.e.m.; *n*=3 mice per time point; unpaired t-test). **c**, Confocal images of whole-mount immunostainings (top view) of murine epidermis at embryonic day 15.5 (E15.5) using anti-Plexin-B1 (green) and anti-Plexin-B2 antibodies (red). Scale bar, 25 μm. **d**, Quantification of K10-positive cells at the indicated embryonic time points (mean ± s.e.m.; *n*=7 mice per genotype and timepoint; unpaired t-test) **e**, TUNEL stainings of murine epidermis at E15.5 and E16.5. Blue: DAPI. Scale bar, 25 μm.

**Supplementary Fig. 2:**
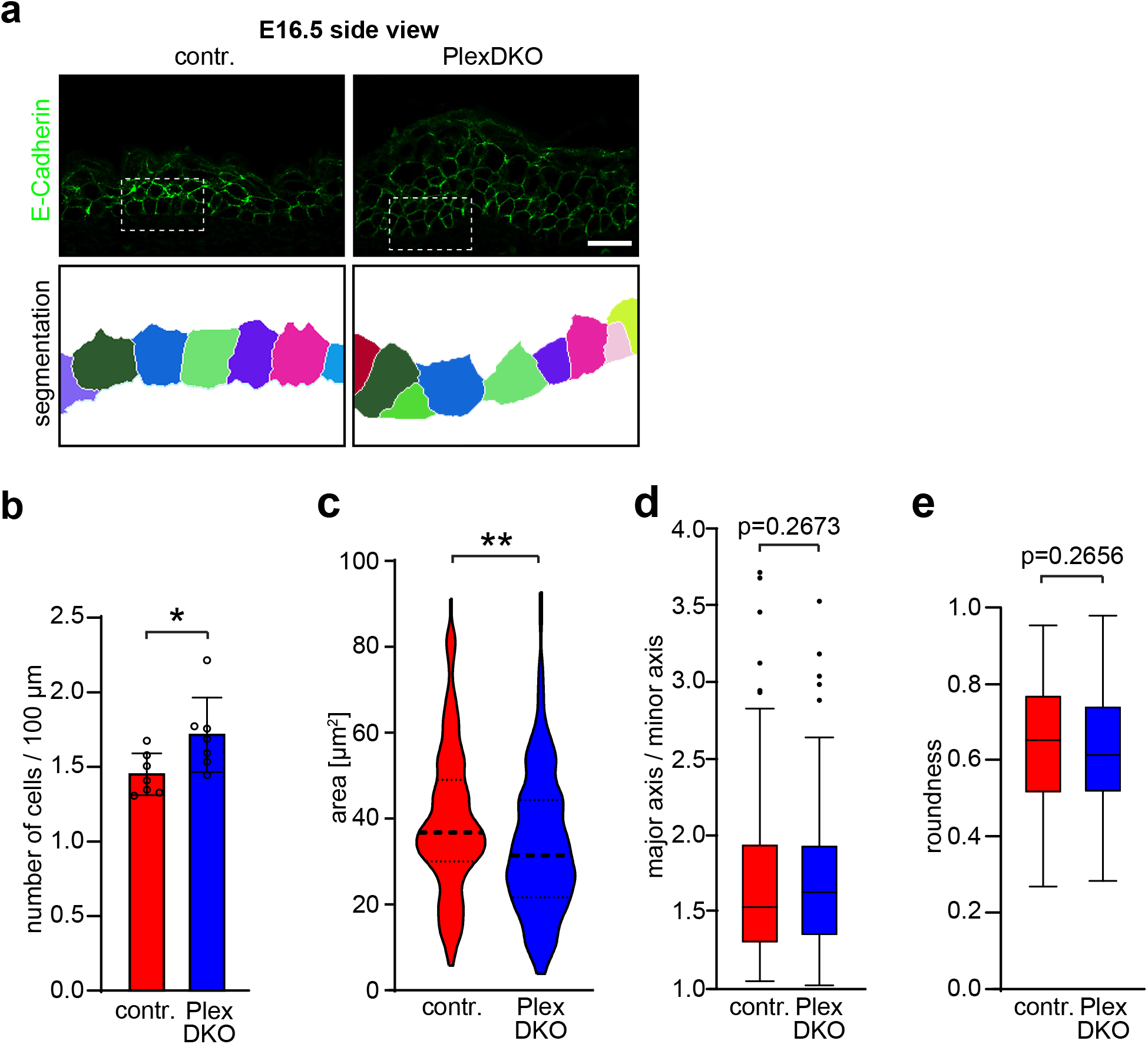
**a**, Upper row: Confocal images of immunostainings (side view) of murine epidermis at E16.5 using an anti-E-cadherin antibody (green). Lower row: Segmentation analyses of the images depicted in the upper row. Scale bar, 25 μm. **b**, Quantification of the number of epidermal stem cells (basal layer) per distance (mean ± s.d.; *n*=7 mice per genotype). **c**, Analysis of epidermal stem cell areas (violin plot with first quartile, median, and third quartile; control: *n*=166 from 7 mice, PlexDKO: *n*=178 from 7 mice; F-test of equality of variances: p=0.6129, Mann-Whitney U test: p=0.0014). **d,e,** Analysis of cell shape anisotropy (box plot with minimum, first quartile, median, third quartile and maximum; control: *n*=166 from 7 mice, PlexDKO: *n*=178 from 7 mice; Mann-Whitney U test).

**Supplementary Fig. 3:**
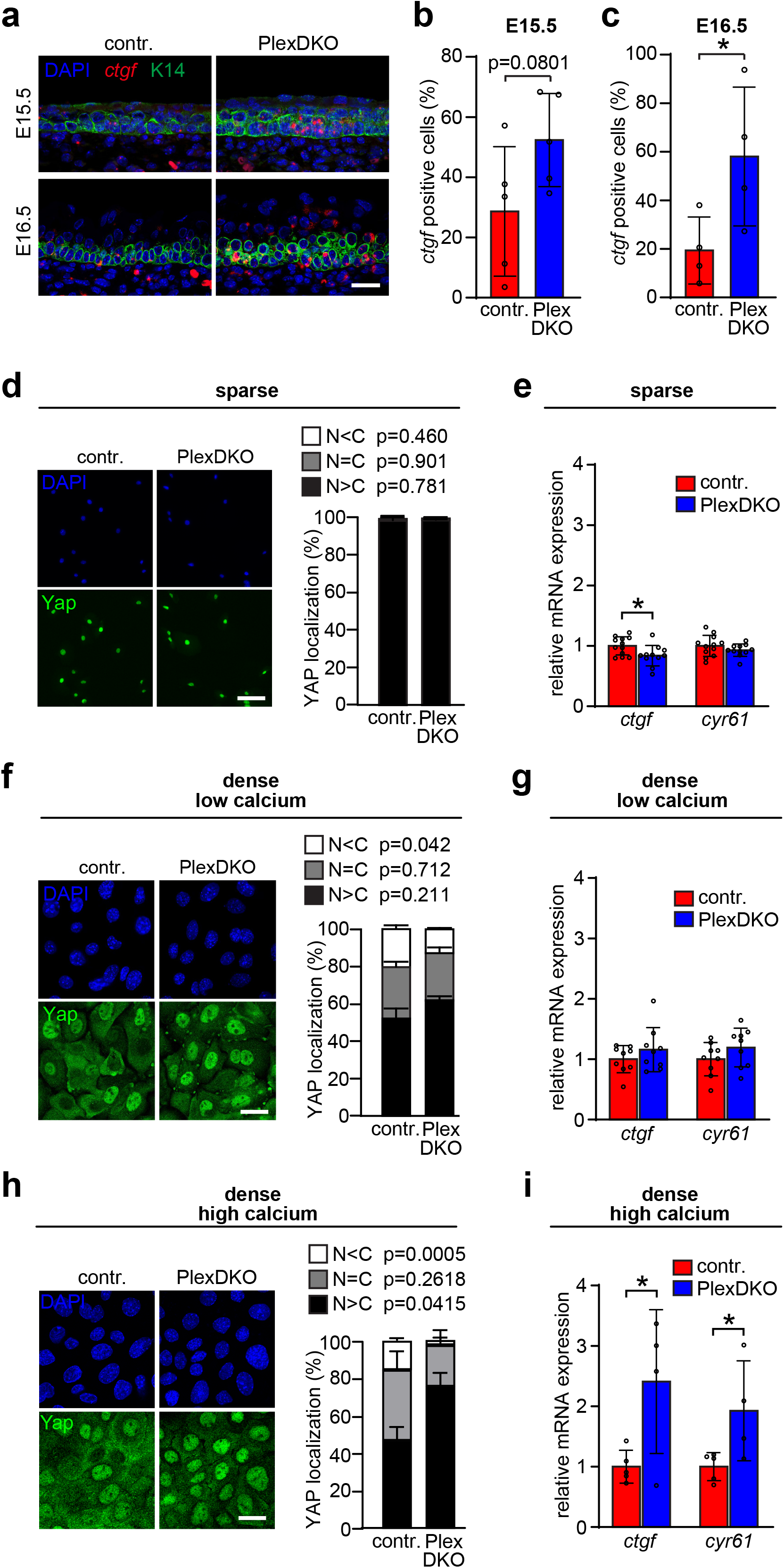
**a**, Confocal images of fluorescent in situ hybridizations for the YAP target gene *ctgf* (red) combined with an immunostaining for K14 (green). Scale bar, 25 μm. **b,c**, Quantification of the data in (a) (mean ± s.d.; E15.5: *n*=5 mice per genotype, E16.5: *n*=4 mice per genotype; unpaired t-test). **d-i**, Primary mouse keratinocytes cultured at different densities and calcium concentrations were immunostained for YAP (green). The left panels show representative images and quantifications of Yap localization. N: nuclear, C: cytoplasmic. Blue: DAPI. The right panels depict mRNA expression levels of the YAP target genes *ctgf* and *cyr61* as determined by quantitative RT-PCR. **d,e,** Primary mouse keratinocytes cultured at low density (“sparse”) and 70 μM Ca^2+^. Quantification of YAP localization: mean ± s.d.; control: *n*=859 from 4 mice, PlexDKO: *n*=1075 from 4 mice; unpaired t-test. Analysis of mRNA expression levels of YAP target genes: mean ± s.d.; control: *n*=12 mice, PlexDKO: *n*=11 mice; unpaired t-test. Scale bar, 100 μm. **f,g,** Primary mouse keratinocytes cultured at high density and 70 μM Ca^2+^. Quantification of YAP localization: mean ± s.d.; control: *n*=468 from 4 mice, PlexDKO: *n*=309 from 3 mice; unpaired t-test. Analysis of mRNA expression levels of YAP target genes: mean ± s.d.; control: *n*=9 mice, PlexDKO: *n*=9 mice; unpaired t-test. Scale bar, 25 μm. **h,i,** Primary mouse keratinocytes cultured at high density and after 3 h of 1.8 mM Ca^2+^. Quantification of YAP localization: mean ± s.d.; control: *n*=923 from 3 mice, PlexDKO: *n*=887 from 3 mice; unpaired t-test. Analysis of mRNA expression levels of YAP target genes: mean ± s.d.; control: *n*=5 mice, PlexDKO: *n*=4 mice; unpaired t-test. Scale bar, 25 μm.

**Supplementary Fig. 4:**
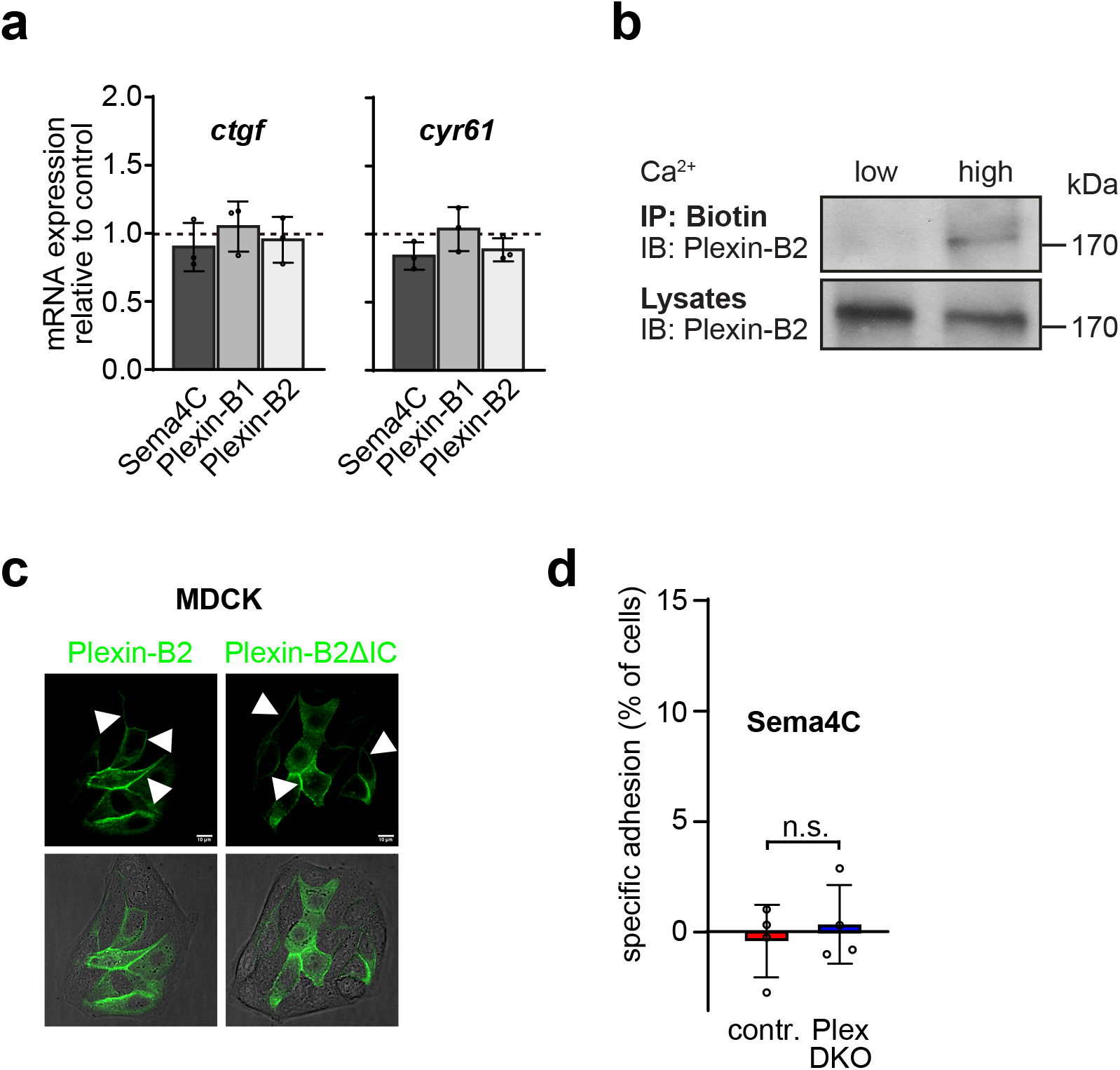
**a**, Primary mouse keratinocytes were incubated without or with recombinant Sema4C, Plexin-B1 or Plexin-B2 (25 nM). After 8 hours, the mRNA expression level of *ctgf* and *cyr61* relative to control (i.e. non-treated cells) was determined by quantitative RT-PCR. **b**, Primary mouse keratinocytes were cultured at low (70 μM) or overnight at high (1.8 mM) Ca^2+^ concentrations. Surface proteins were then biotinylated, precipitated using streptavidin agarose, and surface Plexin-B2 was visualized by Western blotting using an anti-Plexin-B2 antibody. **c**, MDCK renal tubular epithelial cells were engineered to stably express wildtype Plexin-B2 fused to GFP (“Plexin-B2”) or mutant Plexin-B2 lacking the intracellular domain (“Plexin-B2ΔIC”). Shown are representative confocal images. Arrowheads point to cellcell contacts. Scale bars, 10 μm. **d**, 96-well nitrocellulose plates were coated with the recombinant extracellular portion of murine Sema4C, and primary mouse keratinocytes of control or PlexDKO mice were allowed to adhere for 30 min at 37°C. Specific adhesion was quantified as described in Methods.

**Supplementary Fig. 5:**
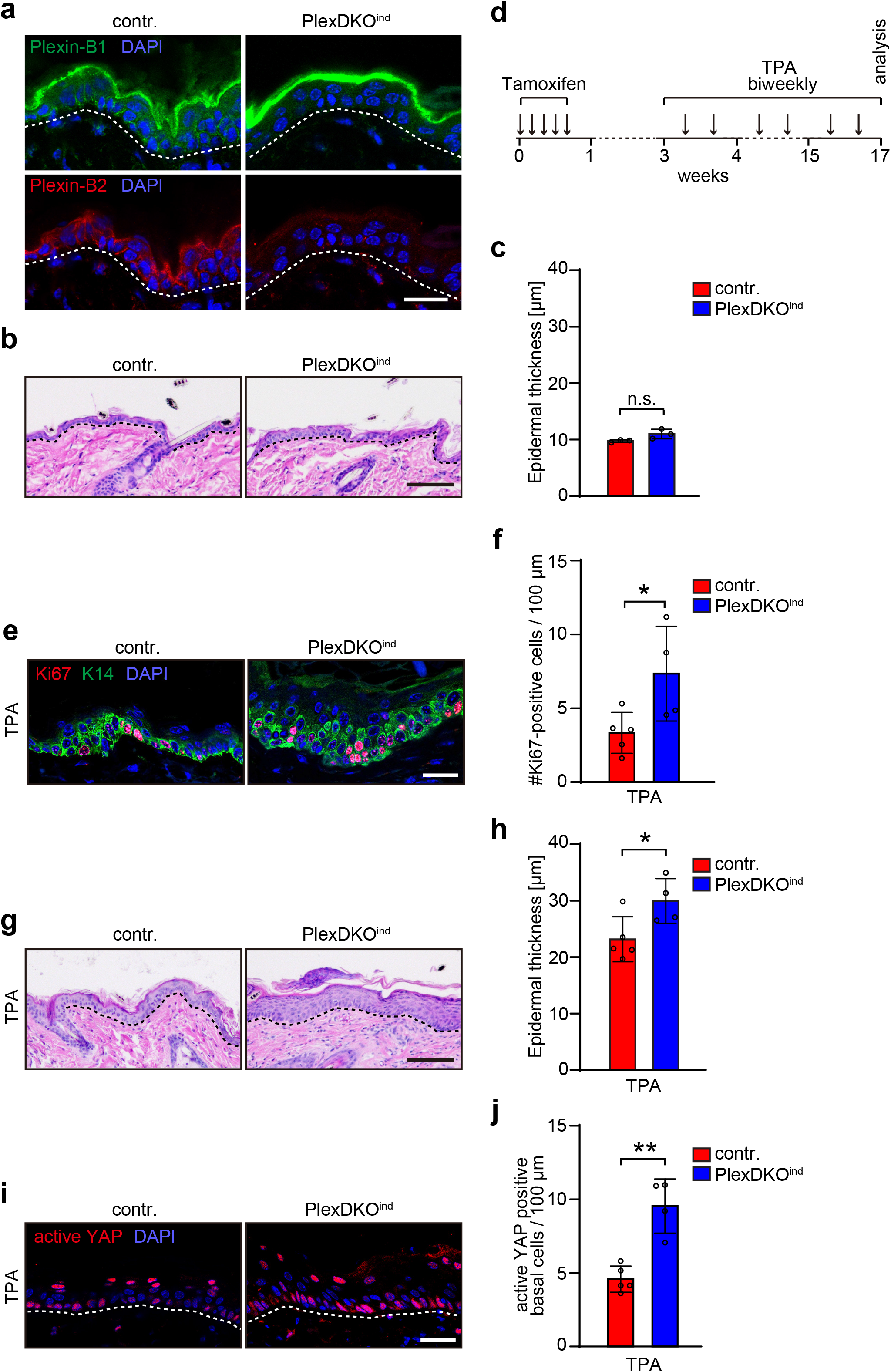
**a**, Adult epidermis-specific tamoxifen-inducible Plexin-B1/Plexin-B2 double-knockout mice (“PlexDKO^ind^”; genotype K14-CreERT;*plxnb1*^flox/flox^;*plxnb2*^flox/flox^) and respective control mice (“contr.”; genotype *plxnb1*^flox/flox^;*plxnb2*^flox/flox^) were treated with tamoxifen for 5 consecutive days, and the epidermis was analyzed. Shown are confocal images of immunostainings using anti-Plexin-B1 (green) or anti-Plexin-B2 (red) antibodies. Scale bar, 25 μm. **b**, H&E stained histological sections of adult skin of mice with the indicated genotypes 17 weeks after tamoxifen treatment. Dashed lines indicate the basement membrane. Scale bar, 100 μm. **c**, Quantification of epidermal thickness (mean ± s.d.; *n*=3 mice per group; unpaired t-test). **d**, Timeline illustrating the treatment regimen of mice with tamoxifen and 12-O-Tetradecanoylphorbol-13-acetate (TPA). **e-j**, Mice with the indicated genotypes were treated according to the scheme depicted in (d). **e**, Confocal images of immunostainings of the skin using anti-Ki67 (red) and anti-K14 antibodies (green). Scale bar, 25 μm. **f**, Quantification of the data in (e) (mean ± s.d.; contr.: *n*=5 mice, PlexDKO: *n*=4 mice; unpaired t-test). **g**, H&E stained histological sections of the skin. Scale bar, 100 μm. **h**, Quantification of epidermal thickness (mean ± s.d.; contr.: *n*=5 mice, PlexDKO: *n*=4 mice; unpaired t-test). **i**, Confocal images of immunostainings of the skin using an anti-active YAP (red) antibody. Scale bar, 25 μm. **j**, Quantification of the data in (i) (mean ± s.d.; contr.: *n*=5 mice, PlexDKO: *n*=4 mice; unpaired t-test).

## REFERENCES

1. Ben Amar, M., Nassoy, P. & LeGoff, L. Physics of growing biological tissues: the complex cross-talk between cell activity, growth and resistance. Philos Trans A Math Phys Eng Sci 377, 20180070 (2019).

2. Irvine, K.D. & Shraiman, B.I. Mechanical control of growth: ideas, facts and challenges. Development 144, 4238–4248 (2017).

3. LeGoff, L. & Lecuit, T. Mechanical Forces and Growth in Animal Tissues. Cold Spring Harb Perspect Biol 8, a019232 (2015).

4. Petridou, N.I. & Heisenberg, C.P. Tissue rheology in embryonic organization. EMBO J 38, e102497 (2019).

5. Petridou, N.I., Spiro, Z. & Heisenberg, C.P. Multiscale force sensing in development. Nat Cell Biol 19, 581–588 (2017).

6. Hannezo, E. & Heisenberg, C.P. Mechanochemical Feedback Loops in Development and Disease. Cell 178, 12–25 (2019).

7. Godard, B.G. & Heisenberg, C.P. Cell division and tissue mechanics. Curr Opin Cell Biol 60, 114–120 (2019).

8. Wickstrom, S.A. & Niessen, C.M. Cell adhesion and mechanics as drivers of tissue organization and differentiation: local cues for large scale organization. Curr Opin Cell Biol 54, 89–97 (2018).

9. Chacon-Martinez, C.A., Koester, J. & Wickstrom, S.A. Signaling in the stem cell niche: regulating cell fate, function and plasticity. Development 145 (2018).

10. Biggs, L.C., Kim, C.S., Miroshnikova, Y.A. & Wickstrom, S.A. Mechanical Forces in the Skin: Roles in Tissue Architecture, Stability, and Function. J Invest Dermatol (2019).

11. Tamagnone, L. et al. Plexins are a large family of receptors for transmembrane, secreted, and GPI-anchored semaphorins in vertebrates. Cell 99, 71–80 (1999).

12. Winberg, M.L. et al. Plexin A is a neuronal semaphorin receptor that controls axon guidance. Cell 95, 903–916 (1998).

13. Worzfeld, T. & Offermanns, S. Semaphorins and plexins as therapeutic targets. Nat Rev Drug Discov 13, 603–621 (2014).

14. Gurrapu, S. & Tamagnone, L. Semaphorins as Regulators of Phenotypic Plasticity and Functional Reprogramming of Cancer Cells. Trends Mol Med 25, 303–314 (2019).

15. Nishide, M. & Kumanogoh, A. The role of semaphorins in immune responses and autoimmune rheumatic diseases. Nat Rev Rheumatol 14, 19–31 (2018).

16. Verlinden, L., Vanderschueren, D. & Verstuyf, A. Semaphorin signaling in bone. Mol Cell Endocrinol 432, 66–74 (2016).

17. Mehta, V. et al. The guidance receptor plexin D1 is a mechanosensor in endothelial cells. Nature 578, 290–295 (2020).

18. Miroshnikova, Y.A. et al. Adhesion forces and cortical tension couple cell proliferation and differentiation to drive epidermal stratification. Nat Cell Biol 20, 69–80 (2018).

19. Zhang, H., Pasolli, H.A. & Fuchs, E. Yes-associated protein (YAP) transcriptional coactivator functions in balancing growth and differentiation in skin. Proc Natl Acad Sci U S A 108, 2270–2275 (2011).

20. Dekoninck, S. et al. Defining the Design Principles of Skin Epidermis Postnatal Growth. Cell (2020).

21. Daviaud, N., Chen, K., Huang, Y., Friedel, R.H. & Zou, H. Impaired cortical neurogenesis in plexin-B1 and -B2 double deletion mutant. Dev Neurobiol 76, 882–899 (2016).

22. Perala, N. et al. Sema4C-Plexin B2 signalling modulates ureteric branching in developing kidney. Differentiation 81, 81–91 (2011).

23. Xia, J. et al. Semaphorin-Plexin Signaling Controls Mitotic Spindle Orientation during Epithelial Morphogenesis and Repair. Dev Cell 33, 299–313 (2015).

24. Hafner, M. et al. Keratin 14 Cre transgenic mice authenticate keratin 14 as an oocyte-expressed protein. Genesis 38, 176–181 (2004).

25. Blanpain, C., Lowry, W.E., Pasolli, H.A. & Fuchs, E. Canonical notch signaling functions as a commitment switch in the epidermal lineage. Genes Dev 20, 3022–3035 (2006).

26. Zhao, B. et al. Inactivation of YAP oncoprotein by the Hippo pathway is involved in cell contact inhibition and tissue growth control. Genes Dev 21, 2747–2761 (2007).

27. Schlegelmilch, K. et al. Yap1 acts downstream of alpha-catenin to control epidermal proliferation. Cell 144, 782–795 (2011).

28. Totaro, A. et al. YAP/TAZ link cell mechanics to Notch signalling to control epidermal stem cell fate. Nat Commun 8, 15206 (2017).

29. Niessen, C.M., Leckband, D. & Yap, A.S. Tissue organization by cadherin adhesion molecules: dynamic molecular and cellular mechanisms of morphogenetic regulation. Physiol Rev 91, 691–731 (2011).

30. Gurrapu, S. et al. Reverse signaling by semaphorin 4C elicits SMAD1/5-and ID1/3-dependent invasive reprogramming in cancer cells. Sci Signal 12 (2019).

31. Sun, T. et al. A reverse signaling pathway downstream of Sema4A controls cell migration via Scrib. J Cell Biol 216, 199–215 (2017).

32. Ohta, K. et al. Plexin: a novel neuronal cell surface molecule that mediates cell adhesion via a homophilic binding mechanism in the presence of calcium ions. Neuron 14, 1189–1199 (1995).

33. Kim, N.G., Koh, E., Chen, X. & Gumbiner, B.M. E-cadherin mediates contact inhibition of proliferation through Hippo signaling-pathway components. Proc Natl Acad Sci U S A 108, 11930–11935 (2011).

34. Lecuit, T. & Yap, A.S. E-cadherin junctions as active mechanical integrators in tissue dynamics. Nat Cell Biol 17, 533–539 (2015).

35. Yap, A.S., Duszyc, K. & Viasnoff, V. Mechanosensing and Mechanotransduction at Cell-Cell Junctions. Cold Spring Harb Perspect Biol 10 (2018).

36. Vasioukhin, V., Bauer, C., Degenstein, L., Wise, B. & Fuchs, E. Hyperproliferation and defects in epithelial polarity upon conditional ablation of alpha-catenin in skin. Cell 104, 605–617 (2001).

37. Noethel, B. et al. Transition of responsive mechanosensitive elements from focal adhesions to adherens junctions on epithelial differentiation. Mol Biol Cell 29, 2317–2325 (2018).

38. Chugh, P. & Paluch, E.K. The actin cortex at a glance. J Cell Sci 131 (2018).

39. Engl, W., Arasi, B., Yap, L.L., Thiery, J.P. & Viasnoff, V. Actin dynamics modulate mechanosensitive immobilization of E-cadherin at adherens junctions. Nat Cell Biol 16, 587–594 (2014).

40. Iglesias-Bartolome, R. et al. Inactivation of a Galpha(s)-PKA tumour suppressor pathway in skin stem cells initiates basal-cell carcinogenesis. Nat Cell Biol 17, 793–803 (2015).

41. Connelly, J.T. et al. Actin and serum response factor transduce physical cues from the microenvironment to regulate epidermal stem cell fate decisions. Nat Cell Biol 12, 711–718 (2010).

42. Ellis, S.J. et al. Distinct modes of cell competition shape mammalian tissue morphogenesis. Nature 569, 497–502 (2019).

43. Watt, F.M., Jordan, P.W. & O’Neill, C.H. Cell shape controls terminal differentiation of human epidermal keratinocytes. Proc Natl Acad Sci U S A 85, 5576–5580 (1988).

44. Mesa, K.R. et al. Homeostatic Epidermal Stem Cell Self-Renewal Is Driven by Local Differentiation. Cell Stem Cell 23, 677–686 e674 (2018).

45. Hanahan, D. & Weinberg, R.A. Hallmarks of cancer: the next generation. Cell 144, 646–674 (2011).

46. Mendonsa, A.M., Na, T.Y. & Gumbiner, B.M. E-cadherin in contact inhibition and cancer. Oncogene 37, 4769–4780 (2018).

47. Kong, Y. et al. Structural Basis for Plexin Activation and Regulation. Neuron 91, 548–560 (2016).

48. Suzuki, K. et al. Structure of the Plexin Ectodomain Bound by Semaphorin-Mimicking Antibodies. PLoS One 11, e0156719 (2016).

49. Douguet, D. & Honore, E. Mammalian Mechanoelectrical Transduction: Structure and Function of Force-Gated Ion Channels. Cell 179, 340–354 (2019).

50. Perala, N.M., Immonen, T. & Sariola, H. The expression of plexins during mouse embryogenesis. Gene Expr Patterns 5, 355–362 (2005).

51. Zielonka, M., Xia, J., Friedel, R.H., Offermanns, S. & Worzfeld, T. A systematic expression analysis implicates Plexin-B2 and its ligand Sema4C in the regulation of the vascular and endocrine system. Exp Cell Res 316, 2477–2486 (2010).

52. Deng, S. et al. Plexin-B2, but not Plexin-B1, critically modulates neuronal migration and patterning of the developing nervous system in vivo. J Neurosci 27, 6333–6347 (2007).

53. Vasioukhin, V., Degenstein, L., Wise, B. & Fuchs, E. The magical touch: genome targeting in epidermal stem cells induced by tamoxifen application to mouse skin. Proc Natl Acad Sci U S A 96, 8551–8556 (1999).

54. Chen, M. et al. Increased glucose tolerance and reduced adiposity in the absence of fasting hypoglycemia in mice with liver-specific Gs alpha deficiency. J Clin Invest 115, 3217–3227 (2005).

55. Yonemura, S., Wada, Y., Watanabe, T., Nagafuchi, A. & Shibata, M. alpha-Catenin as a tension transducer that induces adherens junction development. Nat Cell Biol 12, 533–542 (2010).

56. Rubsam, M. et al. E-cadherin integrates mechanotransduction and EGFR signaling to control junctional tissue polarization and tight junction positioning. Nat Commun 8, 1250 (2017).

57. Grobe, H., Wustenhagen, A., Baarlink, C., Grosse, R. & Grikscheit, K. A Rac1-FMNL2 signaling module affects cell-cell contact formation independent of Cdc42 and membrane protrusions. PLoS One 13, e0194716 (2018).

58. Worzfeld, T. et al. Genetic dissection of plexin signaling in vivo. Proc Natl Acad Sci U S A 111, 2194–2199 (2014).

59. Schindelin, J. et al. Fiji: an open-source platform for biological-image analysis. Nat Methods 9, 676–682 (2012).

60. Faust, U. et al. Cyclic stress at mHz frequencies aligns fibroblasts in direction of zero strain. PLoS One 6, e28963 (2011).

